# Identifying severe COVID-19 risk variants modulating enhancer reporter activity in lung cells

**DOI:** 10.64898/2026.02.26.708234

**Authors:** Giovanna Weykopf, Wendy A Bickmore, Simon C Biddie, Elias T Friman

## Abstract

Common genetic variants contribute to risk for complex human diseases. However, despite thousands of associations, variants modulating disease risk and their functional impact remain largely unknown. This includes SARS-CoV-2 infection, where outcomes range from asymptomatic to fatal. Most host risk variants associated with COVID-19 disease, identified through genome wide association studies, are located in the non-coding genome and may function by altering gene expression in disease-relevant cells and tissues. To address this at scale, we tested >4800 severe COVID-19-associated variants to determine the impact of individual variants and variant combinations on regulatory activity using Self-Transcribing Active Regulatory Region sequencing, a massively-parallel reporter assay, in a lung epithelial cell line (A549). We identify 166 variants within active sequences, of which 29 modulate activity allele-specifically. Evaluating variant combinations, we observe both additive and non-additive effects on regulatory activity. We employ state-of-the-art deep learning models to interpret allele-specific variant effects on regulatory activity and endogenous genomic features. Our work provides a set of prioritised severe COVID-19-associated variants that modulate regulatory activity in lung epithelial cells, candidate transcription factors, and candidate target genes with potential to be disease modifying.

## Introduction

The heterogeneity in disease outcomes following infection by SARS-CoV-2 virus is influenced by pre-existing health conditions, age, and host genetic risk factors^1^. Genome-wide association studies (GWAS) for severe COVID-19 outcomes (involving intensive care admission), have implicated genes involved in viral host entry, lung inflammation, airway mucus defence, and type I interferon response in COVID-19 susceptibility and severity, with severity being highly heritable^2–5^. However, despite the thousands of genetic risk variants identified by GWAS, causal variants and their mechanism of action remain largely unresolved. This is primarily due to linkage disequilibrium (LD), in which non-causal and causal variants are co-inherited. Identifying causal variants, their functional effect, and the biological pathways these perturb can elucidate mechanisms promoting disease progression and inform therapeutic targets. Thus far, efforts to prioritise COVID-19 risk variants have relied largely on computational predictions or functional studies at only one or few loci^6–8^.

Approximately 90% of GWAS variants reside in the non-coding genome^9,10^. Many of these variants likely act by modulating the activity of *cis-*regulatory elements (CREs), including enhancers, that regulate the cell type-or physiological context-specific expression of nearby genes^11,12^. To identify GWAS variants altering enhancer activity in a high-throughput manner, massively-parallel reporter assays (MPRAs), such as Self-Transcribing Active Regulatory Region sequencing (STARR-seq), have been employed^13–18^. STARR-seq measures enhancer activity by quantifying the transcription of self-transcribed candidate sequences which, when active, increase reporter expression beyond basal levels^19^.

Given CREs often function in a tissue-specific, context-dependent, or temporally restricted manner^20–22^, risk variants require assessment in the right context. The main cell types involved in severe COVID-19 pathogenesis are lung epithelial and endothelial cells, and immune cell types^23^. A previous study tested COVID-19 risk variants from two adjacent risk loci, including the gene *LZFTL1*, for regulatory function using MPRA in K562 erythroleukemia cells, prioritising three variants with differential regulatory activity upon SARS-CoV-2 infection^7^. Here, we have used STARR-seq to test 4,894 severe COVID-19-associated risk variants, collated by integrating two GWAS with variants in LD, for their ability to alter enhancer activity in the A549 lung epithelial cell line. Of these, 29 variants displayed allele-specific activity. Additionally, we tested all possible variant combinations which reside in close genomic proximity, finding a further 16 variant pairs with additive or non-additive variant effects. We integrate STARR-seq prioritised variants with datasets indicative of active CREs, and predict the effect on transcription factor (TF) binding and chromatin features. Our work identifies COVID-19 risk variants with regulatory function in lung epithelial cells, demonstrating the value of combining high-throughput allele-specific screening with deep learning models to identify and interpret variant effects and to elucidate disease mechanisms.

## Results

### COVID-19 variant library design

In the lung, SARS-CoV-2 mainly infects type II alveolar epithelial cells, leading to cell death, barrier disruption, and fibrosis in some individuals^23^. To prioritise non-coding COVID-19-associated risk variants functional in the lung epithelium, we screened for risk variants which alter enhancer activity using STARR-seq in the lung epithelial adenocarcinoma cell line A549. A549 cells are derived from alveolar basal epithelial cells of a male patient with non–small cell lung cancer and are widely used as an in vitro model of type II alveolar epithelium^24^. They are amenable to high-throughput screens and allow for the integration of other datasets, including from the Encyclopedia of DNA Elements^25^. We focused on variants identified by the GenOMICC (Genetics Of Mortality In Critical Care) study, given its improved power to detect associations by including only the most severe cases^2,3,26^. We included 2,528 fine-mapped variants from the latest (3^rd^) release ^3^ (**Fig. 1A**), containing functional variants to 99% posterior probability (99% credible set). As no credible set of variants was available for 11/49 independent lead variants, due to multi-ancestry preventing fine-mapping for these regions (see ^3^), we included a further 1,465 variants in LD with the 49 lead variants (r^2^ > 0.7) in European ancestry. As the third release included only common variants (minor allele frequency (MAF) > 0.5%), we additionally included variants from the second GenOMICC release that included rare variants (MAF > 0.02%)^2^, adding 901 further variants. The final set consisted of 4,894 variants, 4,720 of which are single nucleotide polymorphisms (SNPs), with much smaller numbers of small deletions and insertions (**Fig. S1A**). Variants primarily localise to intronic and intergenic regions (**Fig. S1B**), consistent with previous observations across the GWAS catalogue^9^ and are concentrated at a few overrepresented genomic regions (**Fig. S1C**).

**Figure 1.**
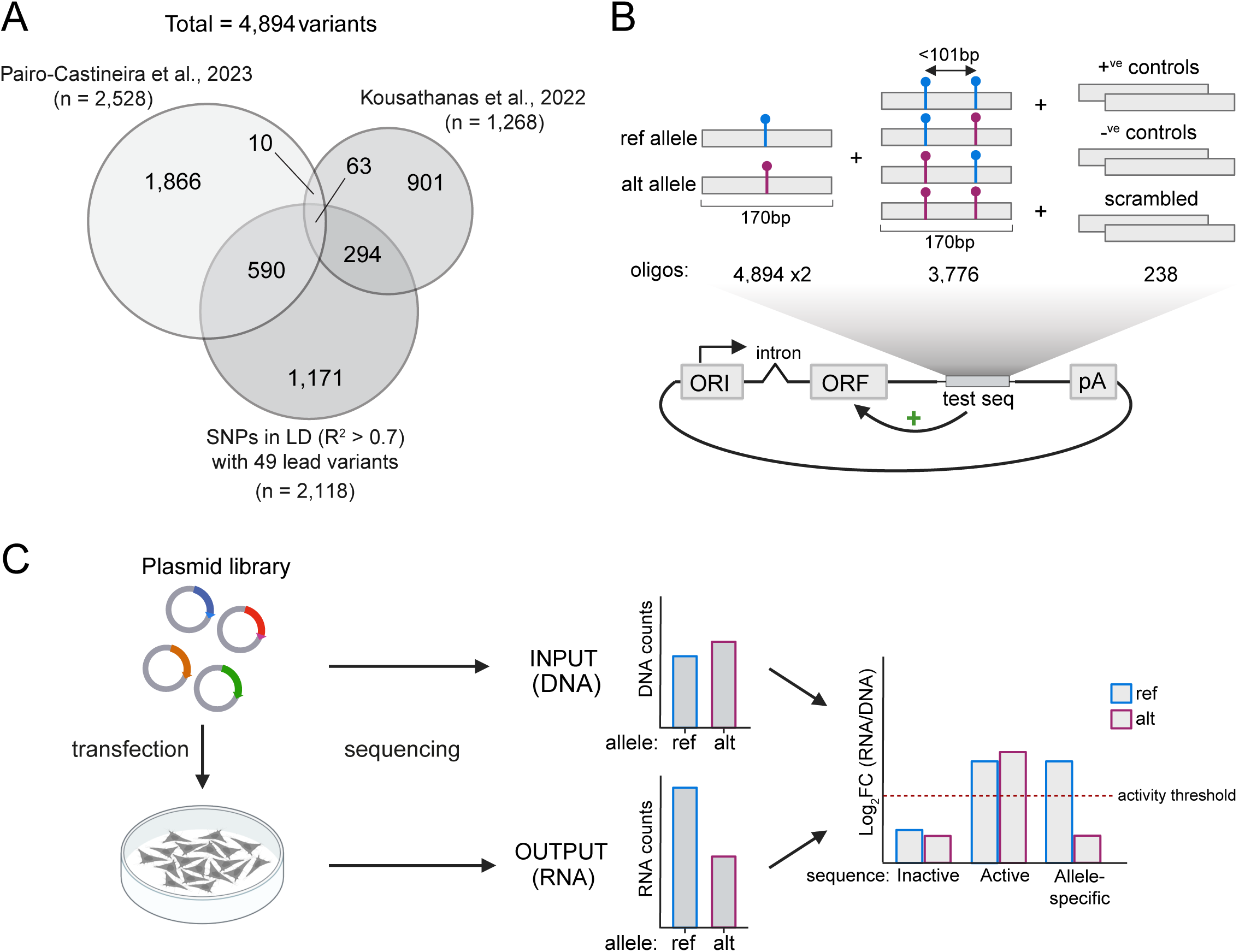
COVID-19 variant STARR-seq library pipeline (A) Venn diagram of variants included in the STARR-seq library and their overlap with each source, including the 99% credible set of fine-mapped variants from Pairo-Castineira et al (2023), the 95% credible set of fine-mapped variants from Kousathanas et al (2022), and variants in LD (r^2^ > 0.7) with any lead variants from Pairo-Castineira et al (2023) using European ancestry. (B) STARR-seq oligonucleotide library design. 170-bp candidate sequences centred on each variant were included as reference (ref) and alternate (alt) alleles (n = 9,790). Additional candidate sequences were included for variants located within 100-bp of at least one other variant in all possible allelic combinations, adding 3,776 further candidate sequences. Up to five variants in proximity were included. 238 170-bp control sequences were included (see Methods). (C) The STARR-seq plasmid library was transfected into A549 cells, input (DNA-) and output (RNA-) sequencing libraries prepared and STARR-seq activity computed as log2FC enrichment of a candidate sequence in RNA normalised to DNA sequencing libraries. Panels (B) and (C) created in BioRender. (https://BioRender.com/3p8cuds).

We designed 170 base pair (bp) oligonucleotides centred on each of the 4,894 risk variants as both reference and alternate allele (two oligonucleotides per variant) (**Fig. 1B**), flanked by 15-bp adapters to facilitate PCR-mediated amplification and cloning. To assess if multiple variants in close proximity alter enhancer function in a combinatorial manner, we additionally tested combinations of variants residing within 100-bp of each other. 777 variants were located in proximity of at least one other variant, the majority of which were variants pairs (n=650) but with up to five variants combined (n=2) (**Fig. S1D**). We designed these to be sequences centred on the middle of the two outermost variants with all possible allelic combinations (**Fig 1B**), adding 3,776 combinatorial oligonucleotides to our STARR-seq library (**Fig S1E, 1C**). As positive controls, we designed 170-bp oligonucleotides centred on the 80 STARR-seq peaks with the highest signal from a genome-wide STARR-seq dataset generated in A549 cells^27^. Negative controls consisted primarily of scrambled sequences of positive controls (see Methods). We cloned this library of 13,802 oligonucleotides into the hSTARR screening vector and performed STARR-seq in five replicates (Fig. 1C)^19,28^.

### Identifying functional COVID-19 variants by STARR-seq in lung epithelial cells

Following sequencing (mean 31 million reads per sample), quality control and alignment, input (DNA) and output (RNA) read counts were generated for each sequence in the library (**Fig. S2A**). 13,461 out of 13,652 sequences (97.5%) passed filtering, and counts correlated highly between replicates (avg. Pearson r DNA = 0.997, RNA = 0.939) (**Fig. S2B**).

To identify sequences with putative enhancer function, we computed the enrichment of normalised RNA over DNA read counts as log_2_ fold-change (log2FC) for each sequence. Candidate sequences were considered active if log2FC was > 1 at a false discovery rate (FDR) < 0.01. Positive (median log2FC = 4.36), but not negative (median log2FC = 0.30), controls were enriched in output (RNA) reads, and classified as active (**Fig. 2A, Fig. S2C,D**). Given that the positive controls were from regions with the highest STARR-seq activity in A549 cells, most COVID-19 variant sequences were expected to show lower activity than the positive controls. Excluding controls, we identified 357 STARR-seq active sequences (2.7% of tested) (**Fig. 2A, B**), 248 of which were single-variant oligonucleotides, corresponding to 166 risk variants for which at least one allele was active (**Fig. 2B**). Variants located within active candidate sequences were on average located closer to the nearest transcription start site (TSS) (median 12 kb) compared to inactive ones (median 16 kb), with only ∼10% within 1 kb of a TSS. (**Fig. S2E**). Similarly, the proportion of active candidate sequences overlapping predicted ENCODE CREs in A549 cells was increased compared to inactive sequences (**Fig. S2F**). In summary, 166 severe COVID-19-associated variants resided in putative CREs in A549 cells.

**Figure 2.**
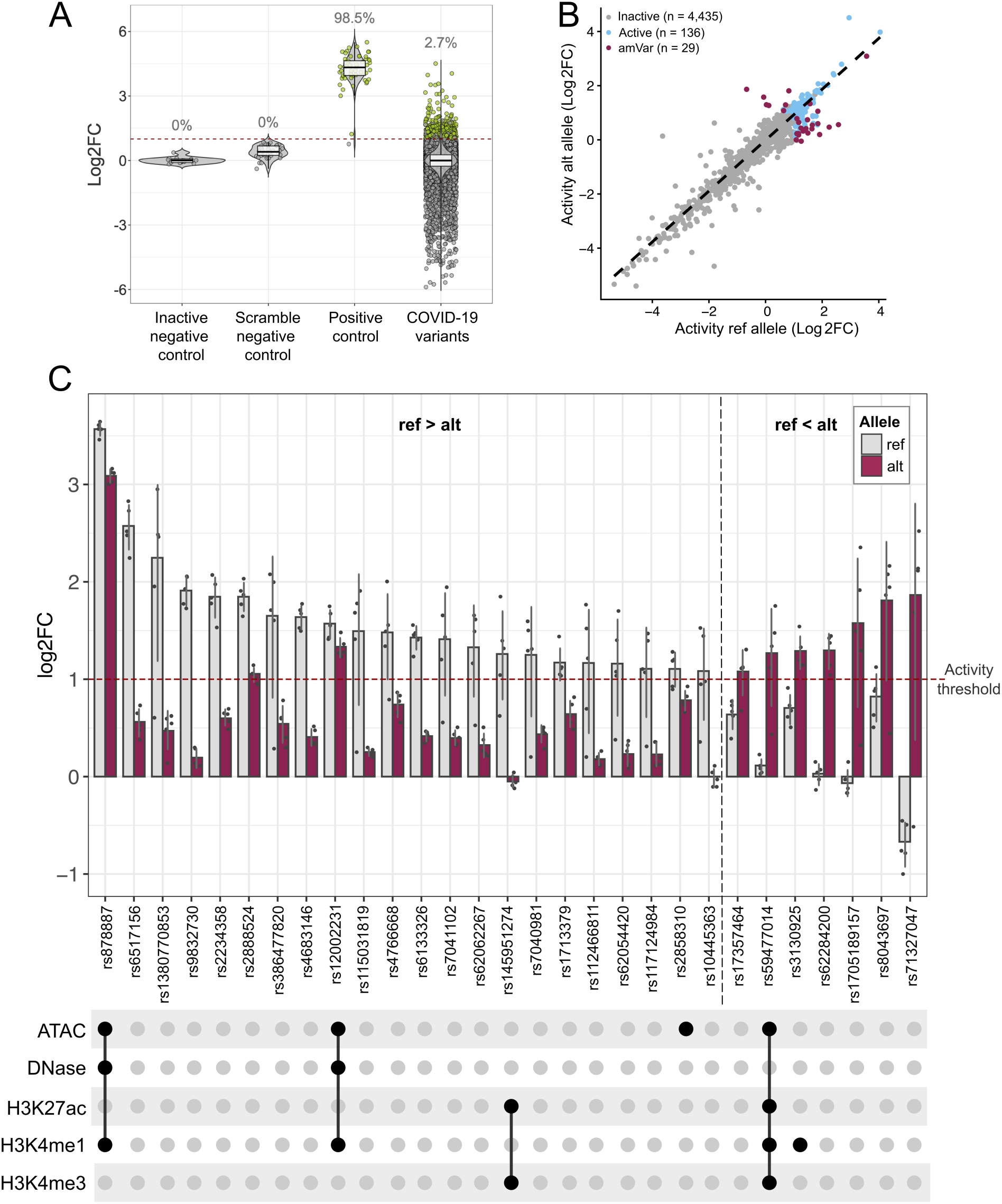
STARR-seq identifies severe COVID-19 risk variants with allele-specific regulatory activity. (A) STARR-seq activity (log2FC) in A549 cells of negative and positive controls, scrambled sequences, and COVID-19 variant sequences across five biological replicates. Sequences are considered active at log2FC > 1 at a false discovery rate < 0.01. Lower and upper hinges of the boxplot correspond to the first and third quartile, respectively. The middle represents the median. (B) Differential STARR-seq activity of alternative (alt) against reference (ref) alleles for each single variant tested as log2FC (n = 4,789). (C) (Top) Enrichment of normalised RNA over DNA read counts for 29 amVars with allele-specific STARR-seq activity showing results for reference and alternative alleles. Allelic differences are significant for all variants shown (FDR < 0.01). Bars represent the mean across five biological replicates; overlayed points represent individual biological replicates. The red line denotes the threshold for active sequences (log2FC > 1). (Bottom) UpSet plot showing the overlap of amVars with A549 ENCODE ATAC-seq, DNase-seq, ChIP-seq for H3K27ac, H3K4me1, and H3K4me3 narrow peaks.

To identify COVID-19 risk variants which have allele-specific STARR-seq activity, we compared normalised RNA and DNA counts for reference and alternate allele. Of 166 variants where at least one allele was active, 29 showed allele-specific differences (FDR<0.01) which we term activity-modulating variants (amVars) (**Fig. 2B, C**). Of those, 22 showed decreased, and 7 increased, activity for the alternate allele compared to the reference allele (**Fig 2C**). These variants are predominantly common, although 4 are rare (minor allele frequency (MAF)<0.005) (**Table 1**). Of those, three are located in intergenic regions of the *IFNA* interferon gene cluster on chromosome 9, a human/ape-specific LINE retrotransposon insertion (**Fig. S3**).

**Table 1.** Features of single variants with allele-specific STARR-seq ac>vity. Table listing each variant as GRCh38 coordinates and rsID with various features. Note for some variants alternative rsIDs exist (not shown). Variant eQTLs in lung from the GTEx Portal (accessed 04/08/2025) showing affected gene(s) with the same direction of effect as observed for STARR-seq, including the normalised effect size (NES) in brackets. Closest protein-coding gene as well as the gene(s) nominated by the GWAS (Pairo-Castineira et al., 2023, Kausathanas et al., 2022) for the risk locus in which the variant was identified are listed. STARR (alt-ref) shows the difference in STARR-seq activity observed for the alternate compared to the reference allele as log2FC, determined using *mpralm*. Allele frequency (allele freq) from gnomAD v.4.1.0 for the alternate alleles averaged across all ancestry groups (Karczewski et al., 2020).

| Variant (hg38) | rsID | Alt allele frequency | Nearest gene | GWAS locus | Putative target genes: (GTex eQTL: NES) in lung | Putative target genes: (GTex sQTL: NES) in lung | STARR-seq effect (alt:ref) |
| --- | --- | --- | --- | --- | --- | --- | --- |
| chr1:155309691:T>G | rs2297480 | 0.2749 | *FDPS* | *EFNA4/TRIM4/THBS3* |  | *FDPS*: 0.58 (sQTL)<br>*FDPS*: -0.61 (sQTL)<br>*FDPS*: -0.36 (sQTL) | 1.135 |
| chr3:45856729:G>A | rs4683146 | 0.4132 | *LZFTL1* | *LZFTL1* |  |  | -1.227 |
| chr3:45947552:T>G | rs2234358 | 0.4858 | *FYCO1/CXCR6* | *LZFTL1* |  | *FYCO1*: -0.52 | -1.238 |
| chr3:45954216:T>G | rs1705189157 | 0.0888 | *FYCO1* | *LZFTL1* |  |  | 1.531 |
| chr3:46256220:A>T | rs9832730 | 0.0698 | *CCR3* | *LZFTL1* |  |  | -1.716 |
| chr3:46269349:T>C | rs71327047 | 0.0512 | *CCR3* | *LZFTL1* |  |  | 2.28 |
| chr3:46326970:T>C | rs2888524 | 0.7357 | *CCR2* | *LZFTL1* | *CCR2*: -0.12 |  | -0.804 |
| chr3:101763892:T>G | rs62284200 | 0.000014 | *CEP97* | *NXPE3* |  | *PCNP*: 0.38<br>*NXPE3*: -0.22 | 1.264 |
| chr6:31198519:C>CG | rs145951274 | 0.1152 | *HCG27* | *CCHCR1* | *MIR6891*: 0.87<br>*TCF19*: 0.23<br>*HLA-C*: 0.38<br>*MICA*: -0.29 | *HLA-C*: -1.2<br>*HLA-B*: -1.2 | -1.265 |
| chr6:31277400:G>C | rs112466811 | 0.1524 | *HLA-C* | *CCHCR1* | *MIR6891*: 0.80<br>*HLA-C*: 0.41<br>*TCF19*: 0.19<br>*MICA*: -0.26<br>*HLA-B*: -1.8 | *HLA-C*: -1.2<br>*HLA-B*: -1.2 | -0.895 |
| chr6:31277403:T>C | rs115031819 | 0.1524 | *HLA-C* | *CCHCR1* | *MIR6891*: 0.80<br>*HLA-C*: 0.41<br>*TCF19*: 0.19<br>*MICA*: -0.26<br>*HLA-B*: -1.8 | *HLA-C*: -1.2<br>*HLA-B*: -1.2 | -1.148 |
| chr6:31495844:A>G | rs3130925 | 0.8035 | *MICB* | *FOXP4* | *HLA-S*: 0.49<br>*HLA-C*: -0.27<br>*HCG27*: 0.20 | *HLA-C*: 1.0<br>*DDX39B*: 1.0<br>*HLA-B*: 1.0 | 0.573 |
| chr6:32700546:G>A | rs2858310 | 0.6446 | *HLA-DQA2/DQB1* | *HLADQA1* | *HLA-DQA2*: 0.69<br>*HLA-DQB1*: -0.46<br>*HLA-DRB1*: -0.35 | *HLA-DRB1*: 0.76, -0.78<br>*HLA-DRB5/6*: 0.76, -0.78 | -0.32 |
| chr8:60487028:C>T | rs6471885 | 0.4312 | *RAB2A* | *RAB2A* | *RAB2A*: 0.14 |  | 0.434 |
| chr9:21211717:C>T | rs7041102 | 0.0368 | *IFNA10/IFNA16* | *IFNA10* |  |  | -0.942 |
| chr9:21211718:A>G | rs7040981 | 0.0369 | *IFNA10/IFNA16* | *IFNA10* |  |  | -0.737 |
| chr9:21294504:G>A | rs12002231 | 0.0346 | *IFNA5* | *IFNA10* |  |  | -0.24 |
| chr10:79518244:G>A | rs1713379 | 0.4164 | *EIF5AL1* | *SFTPD* | *NUTM2B*: -0.44<br>*BEND3P3*: -0.29 | *SFTPA1*: 0.44, -0.29<br>*SFTPA2*: -0.20 | -0.513 |
| chr12:112926975:T>C | rs4766668 | 0.7432 | *OAS1/OAS3* | *OAS1* |  | *OAS1*: -1.5 | -0.703 |
| chr16:89202660:C>G | rs8043697 | 0.1991 | *SLC22A31* | *SLC22A31* | *ZNF778*: 0.2<br>*CDH15*: 0.29 | *ACSF3*: -0.8<br>*SLC22A31*: -0.48 | 0.89 |
| chr17:45739289:C>G | rs62054420 | 0.1499 | *CRHR1* | *KANSL1* |  | *KANSL1*: -1.6 | -0.902 |
| chr17:45835216:C>T | rs878887 | 0.1573 | *CRHR1* | *KANSL1* |  | *KANSL1*: -1.6 | -0.475 |
| chr17:45837192:G>A | rs10445363 | 0.1421 | *CRHR1* | *KANSL1* | *LRRC37A2*: 0.95<br>*LRRC37A*: 0.89<br>*MAPT*: 0.59 | *KANSL1*: -1.6<br>*PLEKHM1*: -0.56<br>*ARHGAP27*: -0.54 | -1.063 |
| chr17:45974222:C>G | rs117124984 | 0.1611 | *MAPT* | *KANSL1* | *LRRC37A2*: 0.95<br>*LRRC37A*: 0.89<br>*MAPT*: 0.58 | *KANSL1*: -1.6<br>*PLEKHM1*: -0.55<br>*ARHGAP27*: -0.55 | -0.832 |
| chr17:46013488:C>T | rs62062267 | 0.1431 | *MAPT* | *KANSL1* | *LRRC37A2*: 0.96<br>*LRRC37A*: 0.90<br>*MAPT*: 0.59 | *KANSL1*: -1.7 | -0.968 |
| chr19:48704000:T>C | rs603985 | 0.5018 | *FUT2* | *FUT2* | *IZUMO1*: 0.21<br>*FUT2*: -0.16 |  | -0.997 |
| chr19:48708231:C>A | rs1380770853 | 0.02472 | *FUT2* | *FUT2* |  |  | -1.631 |
| chr20:6507080:G>A | rs6133326 | 0.4285 | *BMP2* | *CASC20* |  |  | -1.01 |
| chr21:33261740:C>G | rs6517156 | 0.3615 | *IFNAR2* | *IFNAR2/IL10RB* | *IFNAR2*: -0.24 | *IFNAR2*: 0.40 | -2.003 |

STARR-seq is an episomal assay, so it does not assess endogenous enhancer function. To evaluate which putative CREs harbouring amVars may be endogenously active in A549 cells, we intersected amVars with A549 ATAC-seq, DNase-seq, and ChIP-seq datasets for H3K27ac, H3K4me1, and H3K3me3 from ENCODE^25^, all of which are associated with endogenous CRE activity. Of 29 variants, six overlapped peaks from at least one dataset (**Fig 2C**). Of those, rs145951274, rs3130925, and rs2858310 reside within the human leukocyte antigen (HLA) region on chromosome 6, including at least 132 protein-coding genes encoding for MHC class I, II and III complexes^29^.

### Variant pairs alter activity individually, additively or non-additively

Multiple sequence changes can alter an enhancer’s activity^30,31^. It is therefore plausible that combinations of variants in close proximity and high LD can underlie the GWAS association of a locus^32,33^. To assess this in the context of severe COVID-19 risk variants, our library included combinations of 777 variants which reside within 100-bp of at least one other variant as additional candidate sequences (**Fig. 1B**). Of 3,776 combinatorial sequences tested, 48 variant combinations had at least one allelic combination that showed STARR-seq activity (log2FC>1, FDR<0.01) (**Fig, S4A**). We subsequently focused on sequences with higher STARR-seq activity (log2FC > 1.5) of at least one allele, resulting in 16 combinations, all of which were variant pairs (**Fig. 3A**, **Table 2**). The activity of prioritised combinatorial oligonucleotides correlated highly with the activity of the respective single variant oligonucleotides matched in genotype (Pearson r= 0.802) (**Fig S4B**), showing that STARR-seq activity is largely insensitive to slight shifts in sequence.

**Fig 3.**
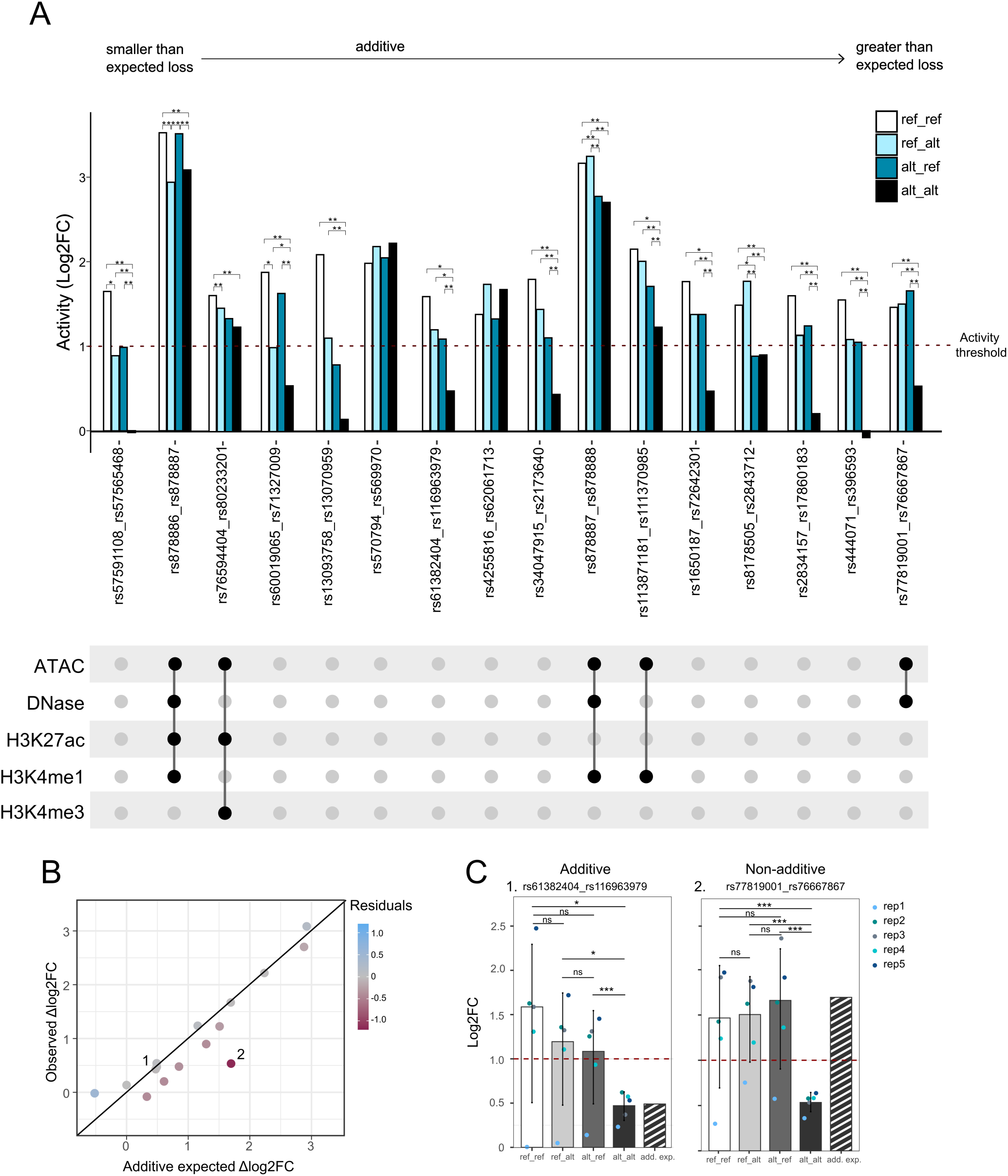
Variant combinations can have additive or non-additive effects on STARR-seq activity. (A) Top: STARR-seq activity (log2FC) of combinatorial oligos for 16 variant pairs where at least one allele shows log2FC > 1.5 at FDR < 0.01, for each variant combination. STARR-seq activity threshold (log2FC >1) indicated as dotted red line. Pairwise significance computed by *mpralm* indicated where padj < 0.05 (*) or padj < 0.01 (**) between alleles. Bottom: UpSet plot showing the overlap of genomic intervals spanning from variant 1 to variant 2 with A549 ENCODE ATAC-seq, DNase-seq, and H3K27ac, H3K4me1, and H3K4me3 ChIP-seq narrow peaks. (B) Observed vs expected STARR-seq Δlog2FC (log2FC alt - log2FC ref) for alt_alt alleles if both variant effects interacted additively. Colour indicates the difference between observed and expected effects (Residuals). Numbers 1 and 2 indicate examples highlighted in (C). (C) Observed STARR-seq activity for all four alleles and additive expected STARR-seq activity for the alt_alt allele for variant pair (left); rs61382404 and rs116963979 and (right); rs77819001 and rs76667867. Bars indicate activity averaged across five biological replicates, points show individual biological replicates. ns = not significant, * < 0.05, ** < 0.01 and *** < 0.001.

**Table 2.**
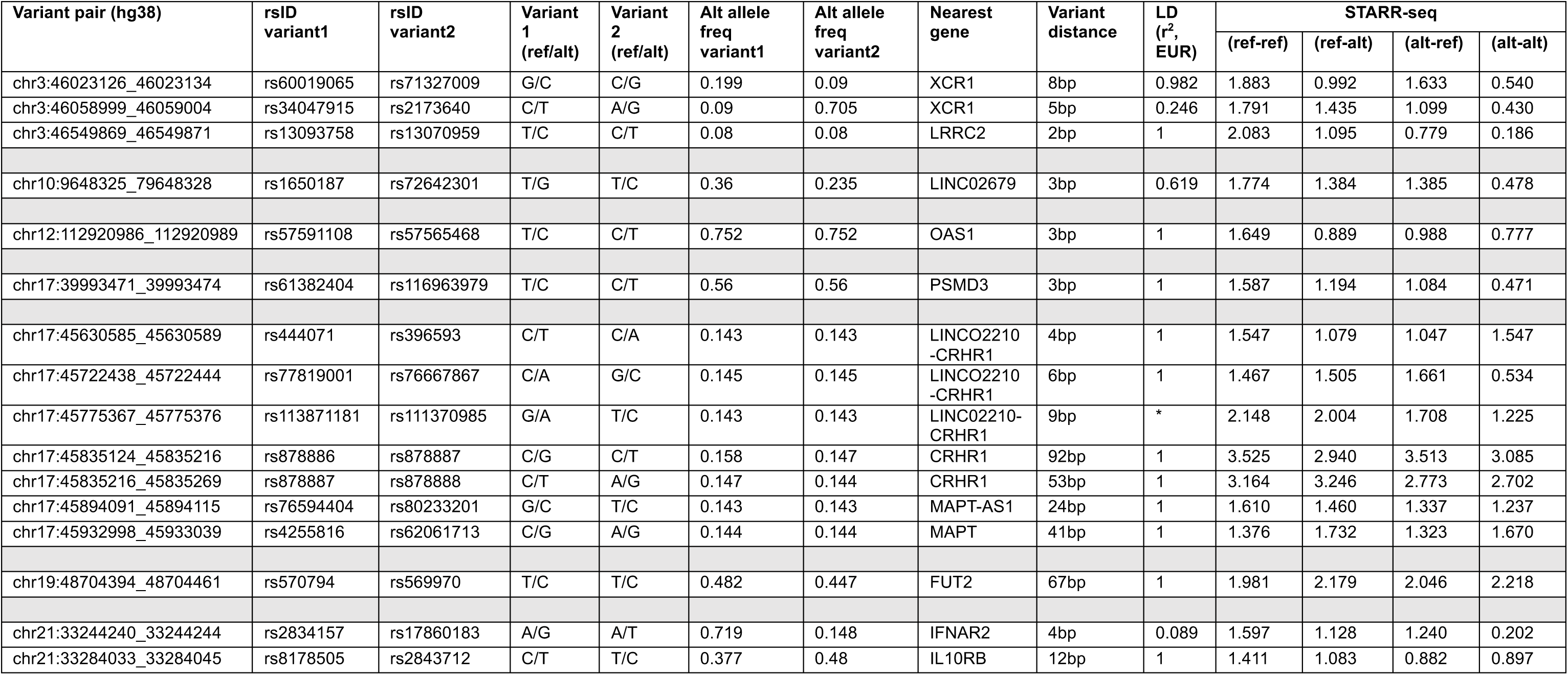
Features of variant pairs with STARR-seq activity. Table listing each variant pair as GRCh38 coordinates (chr:variant1_variant2) and rsID with various features. mAllele frequency (allele freq) from gnomAD v.4.1.0 for the alternate alleles averaged across all ancestry groups (Karczewski et al., 2020). Linkage disequilibrium (LD) in European ancestry calculated using the LDpair tool from LDlink (Machiela and Chanock 2015) (hSps://ldlink.nih.gov/ldpair). *No LD value available as one variant is not in the 1000 Genomes reference panel. STARR-seq ac)vity for all possible allelic combinations.

| Variant pair (hg38) | rsID<br>variant1 | rsID<br>variant2 | Variant<br>1<br>(ref/alt) | Variant<br>2<br>(ref/alt) | Alt allele<br>freq<br>variant1 | Alt allele<br>freq<br>variant2 | Nearest<br>gene | Variant<br>distance | LD<br>(r<sup>2</sup>,<br>EUR) | STARR-seq |  |  |  |
| --- | --- | --- | --- | --- | --- | --- | --- | --- | --- | --- | --- | --- | --- |
|  |  |  |  |  |  |  |  |  |  | (ref-ref) | (ref-alt) | (alt-ref) | (alt-alt) |
| chr3:46023126_46023134 | rs60019065 | rs71327009 | G/C | C/G | 0.199 | 0.09 | XCR1 | 8bp | 0.982 | 1.883 | 0.992 | 1.633 | 0.540 |
| chr3:46058999_46059004 | rs34047915 | rs2173640 | C/T | A/G | 0.09 | 0.705 | XCR1 | 5bp | 0.246 | 1.791 | 1.435 | 1.099 | 0.430 |
| chr3:46549869_46549871 | rs13093758 | rs13070959 | T/C | C/T | 0.08 | 0.08 | LRRC2 | 2bp | 1 | 2.083 | 1.095 | 0.779 | 0.186 |
| chr10:9648325_79648328 | rs1650187 | rs72642301 | T/G | T/C | 0.36 | 0.235 | LINC02679 | 3bp | 0.619 | 1.774 | 1.384 | 1.385 | 0.478 |
| chr12:112920986_112920989 | rs57591108 | rs57565468 | T/C | C/T | 0.752 | 0.752 | OAS1 | 3bp | 1 | 1.649 | 0.889 | 0.988 | 0.777 |
| chr17:39993471_39993474 | rs61382404 | rs116963979 | T/C | C/T | 0.56 | 0.56 | PSMD3 | 3bp | 1 | 1.587 | 1.194 | 1.084 | 0.471 |
| chr17:45630585_45630589 | rs444071 | rs396593 | C/T | C/A | 0.143 | 0.143 | LINC02210<br>-CRHR1 | 4bp | 1 | 1.547 | 1.079 | 1.047 | 1.547 |
| chr17:45722438_45722444 | rs77819001 | rs76667867 | C/A | G/C | 0.145 | 0.145 | LINC02210<br>-CRHR1 | 6bp | 1 | 1.467 | 1.505 | 1.661 | 0.534 |
| chr17:45775367_45775376 | rs113871181 | rs111370985 | G/A | T/C | 0.143 | 0.143 | LINC02210-CRHR1 | 9bp | * | 2.148 | 2.004 | 1.708 | 1.225 |
| chr17:45835124_45835216 | rs878886 | rs878887 | C/G | C/T | 0.158 | 0.147 | CRHR1 | 92bp | 1 | 3.525 | 2.940 | 3.513 | 3.085 |
| chr17:45835216_45835269 | rs878887 | rs878888 | C/T | A/G | 0.147 | 0.144 | CRHR1 | 53bp | 1 | 3.164 | 3.246 | 2.773 | 2.702 |
| chr17:45894091_45894115 | rs76594404 | rs80233201 | G/C | T/C | 0.143 | 0.143 | MAPT-AS1 | 24bp | 1 | 1.610 | 1.460 | 1.337 | 1.237 |
| chr17:45932998_45933039 | rs4255816 | rs62061713 | C/G | A/G | 0.144 | 0.144 | MAPT | 41bp | 1 | 1.376 | 1.732 | 1.323 | 1.670 |
| chr19:48704394_48704461 | rs570794 | rs569970 | T/C | T/C | 0.482 | 0.447 | FUT2 | 67bp | 1 | 1.981 | 2.179 | 2.046 | 2.218 |
| chr21:33244240_33244244 | rs2834157 | rs17860183 | A/G | A/T | 0.719 | 0.148 | IFNAR2 | 4bp | 0.089 | 1.597 | 1.128 | 1.240 | 0.202 |
| chr21:33284033_33284045 | rs8178505 | rs2843712 | C/T | T/C | 0.377 | 0.48 | IL10RB | 12bp | 1 | 1.411 | 1.083 | 0.882 | 0.897 |

Five of the 16 variant combinations resided within accessible chromatin (ATAC-seq) and either DNase-seq, H3K27ac, H3K4me1, or H3K4me3 ChIP-seq peaks from A549 cells (**Fig. 3A**), suggestive of endogenously active enhancers. Interestingly, all five of these variants are located at the same GWAS risk locus, a region on chromosome 17 encompassing multiple plausible target genes (**Fig. S5**). Two STARR-seq active prioritised variant pairs (rs77819001; rs76667867 and rs113871181;rs111370985) are located 50kb and 400bp, respectively, upstream of the TSS of *Corticotropin Releasing Hormone Receptor 1* (*CRHR1)*. Two further variant pairs (rs878886;rs878887 and rs878887;rs878888) are located within the *CRHR1* 3’ untranslated region (UTR) (**Fig. S5**). The remaining variant pair (rs76594404;rs80233201) resides 500bp upstream of the *MAPT* promoter. Alt-alt alleles are associated with increased expression of a long non-coding RNA *LINC02210* and decreased *KANSL1* mRNA splicing in lung (**Fig. S5**).

For variant pairs within active candidate sequences, either variant or both in combination can modulate STARR-seq activity (**Fig 3A**). To determine if variant effects are additive, we calculated the expected STARR-seq activity in the presence of both variants (alt_alt allele) based on either variant alone (ref_alt and alt_ref alleles). We found that 9/16 (56%) variant pairs interacted approximately additively, i.e. the expected log2FC closely resembled that observed (**Fig. 3B**), for example rs61382404 and rs116963979 which reside 3bp apart (**Fig. 3C, Table S1**). An additive model better captured the observed combined STARR-seq activity overall, than a multiplicative model (**Fig. S4C,D, Table S1**).

Interestingly, a subset of variant pairs was not well captured by either the additive or multiplicative model (here called non-additive) (**Fig 3B,C**). We highlight rs77819001 and rs76667867 as an example, 6bp apart, where only in the presence of both variants, STARR-seq activity is lost. We conclude that proximal variants combine mainly additively, but sometimes non-additively, showing that interdependent effects can be missed by studying variants in isolation.

### Implementation of deep learning can aid the interpretation of allele-specific effects

Determining the effect of variants on TF binding and other functional outcomes is not trivial. To evaluate the ability of state-of-the-art deep learning models to predict our experimentally validated amVars, and their usefulness in providing additional information about the variants, we used two complementary models. We first used AlphaGenome ^34^, trained on readouts from multiple genomic data modalities and cell types including chromatin accessibility, histone modifications, gene expression, and TF binding in A549 cells, to predict the regulatory impact of variants within STARR-seq active sequences (n = 166). Filtering for meaningful predictions based on quantile scores (absolute value>0.99), 18/29 (62.1%) amVars and 64/137 (46.7%) non-amVars within STARR-seq active sequences were predicted to significantly alter at least one feature (**Fig. 4A, Fig. S6A-B**). Changes to RNA-seq were most frequently predicted (n=155) (**Fig. 4B, Fig. S6A**) but with negligible effect sizes (median raw score = - 0.0069) (**Figs. 4C, S6B**) which showed negligible correlation with amVar effects on STARR-seq activity (Pearson r =-0.17, Spearman’s p =-0.057, AUC = 0.52) (**Fig. 4C, Fig S6C**). TF ChIP-seq predictions correlated poorly in aggregate with STARR-seq observed amVar effects (Δlog2FC) (Pearson r = 0.23, Spearman’s p = 0.113, n = 70) (**Fig. 4C**), likely as this includes predictions for multiple TFs. We therefore did not consider TF ChIP-seq and RNA-seq predictions further. In contrast, ATAC-seq (n = 4), histone modification ChIP-seq including H3K27ac, H3K4me1, and H3K4me3 (n = 17), DNase-seq (n = 5) and CAGE-seq predictions (n = 5) correlated positively with amVars and less or negatively with non-amVars (**Fig. 4C**). While only 9/29 amVars (31%) were predicted to impact any of these features (**Fig. 4B**), the predicted direction of effect agreed with the STARR-seq observed impact (i.e. loss or gain) in all instances (**Fig. 4C**, **Table S2**). Overall, AlphaGenome correctly predicted a subset of amVars (area under curve (AUC) = 0.68-0.75) (**Fig S6C**), however, many true allele-specific effects were missed.

**Fig 4.**
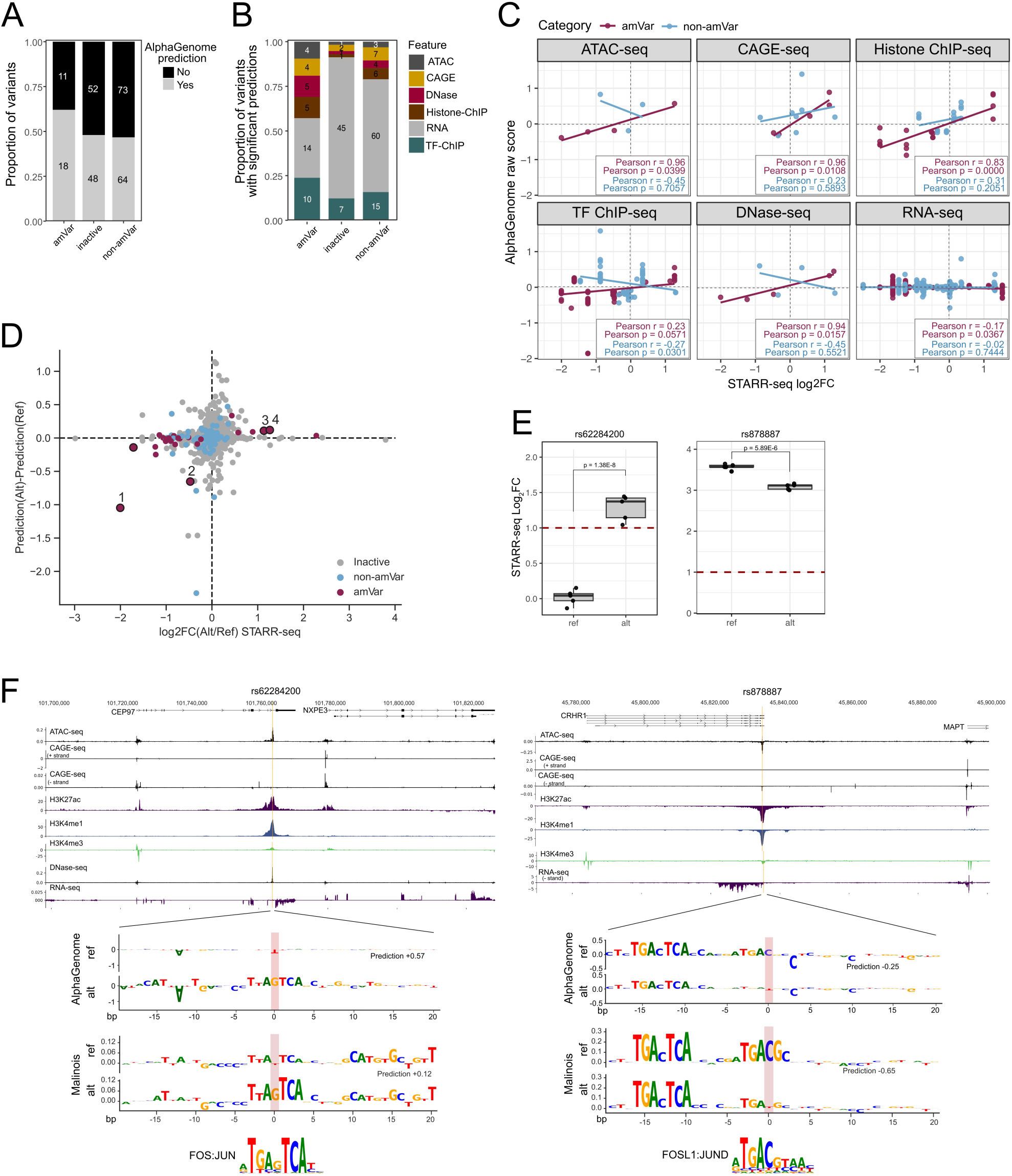
Deep learning models interpret observed allele-specific variant effects. (A) Proportion of amVars, and variants within inactive or active sequences without allelic effects (non-amVars) for which AlphaGenome generated predictions with an absolute quantile score >0.99 for any feature. Absolute number of variants for each category labelled. (B) Proportion of AlphaGenome predictions for each feature for amVars, inactive variants and non-amVars (active, not allele-specific) that were predicted to have any allele-specific effects, as displayed in (A). Note variants can have predictions across multiple features, the absolute number of predictions per category is indicated. (B) Correlation between AlphaGenome predicted change (raw score) and STARR-seq observed allelic effect (log2FC) for each feature predicted by AlphaGenome for amVars and non-amVars. (C) Difference between Malinois MPRA activity prediction for A549 cells between alternative and reference allele compared to observed STARR-seq allelic difference (log2FC alt - log2FC ref). Numbered points correspond to amVars with a Malinois prediction score >=0.5 and are (1) rs6517156, (2) rs878887, (3) rs2297480, and (4) rs62284200. (D) STARR-seq activity for reference and alternative allele of (left) rs62284200 (right) rs878887. Boxplot showing the median, second and third quantiles averaged across five biological replicates displayed as individual points. Dotted red line indicates the STARR-seq activity threshold (log2FC > 1). (E) AlphaGenome alternative-reference predicted genome tracks for (left) rs62284200 and (right) rs878887 within 100kb centred on the variant in A549 cells. The difference in predictions between alternative and reference alleles for features with an absolute quantile score > 0.99 is shown. Below this, the AlphaGenome (top panel) and Malinois (bottom panel) contribution scores for reference and alternative alleles of rs62284200 (ATAC-seq) or rs878887 (CAGE-seq) and MPRA activity predictions, respectively. The matched JASPAR motifs for FOS:JUN (MA0099.3, rs62284200, left) and FOSL1:JUND (MA1143.1, rs878887, right) are at the bottom.

As a complementary approach, we predicted the A549-specific MPRA activity of our library sequences using Malinois, a task-specific deep convolutional neural network model^35^. Malinois predicted most COVID-19 candidate sequences to be inactive (**Fig. S6D**), in agreement with our experimental observations (**Fig. 2A**), and distinguished between positive and negative controls (**Fig. S6D**). Higher activity was predicted for STARR-seq active compared to inactive sequences overall (**Fig. S6E**), although over 70% of active sequences were predicted to have low activity comparable to that of negative controls (**Fig. S6D-E**). We found negligible differences in predictions for most amVars and large differences for some non-amVars, but with reasonable overall performance (AUC = 0.73) (**Fig. 4D, Fig. S6F-G**). Focusing on amVars where either allele has a predicted activity score of at least 0.5 (7/29), the predicted direction of effect was correct in all instances (**Fig. 4D, highlighted points**). Overall, where Malinois and AlphaGenome generated differential predictions for amVars, loss or gain was predicted with perfect accuracy, albeit with a low number of tested variants (**Fig. 4C,D**). However, both models exhibited a high false negative rate for amVars, indicating they are not well suited to capture allele-specific effects in weak enhancers as observed in our library.

Lastly, we assessed whether AlphaGenome and Malinois can provide additional information on observed amVar effects to aid hypothesis generation by *in-silico* mutagenesis (ISM) of correctly predicted variants and matching of identified sequence patterns to known TF motifs. rs62284200, residing in the last intron of *Centrosomal Protein 97* (*CEP97*) and 15kb upstream of the TSS of *NPEX3* (**Fig. S7A**), caused gain of STARR-seq activity (**Fig. 4E**) and predicted increase of DNase-seq, ATAC-seq, H3K27ac, and H3K4me1 ChIP-seq signals (**Fig. 4F**). The alternative allele creates a FOS:JUN motif (**Fig. 4F)** positively contributing to MPRA activity (+0.12) and ATAC-seq predictions (+0.57). FOS/JUN family heterodimers are pioneer TFs known to recruit chromatin remodelling complexes^36^; therefore, it is plausible that rs62284200 drives activity of a regulatory element otherwise inactive in A549 cells.

Conversely, rs878887, located in the 3’ UTR of *CRHR1* (**Fig. S5A**) and causing decreased STARR-seq signal (**Figs. 2C, 4E**), is predicted to decrease chromatin accessibility, CAGE-seq signal, H3K27ac, and H3K4me1 ChIP-seq signals in A549 cells (**Fig. 4F**) and is an sQTL for decreased *KANSL1* intron exclusion (**Fig. S5C**). The variant disrupts a FOSL1:JUND motif, negatively contributing to MPRA activity (-0.65) and CAGE-seq (-0.25) predictions (**Fig. 4F**). The closely related TF FOSL2, binding to a highly similar motif, is detected by ChIP-seq at the variant in A549 cells (**Fig S5A**).

rs6517156 in an intron of *IFNAR2*, causing the most pronounced loss of STARR-seq activity observed for any amVar (**Figs. S7B, 2C**), is predicted to decrease ATAC-seq, H3K27ac and H3K4me1 ChIP-seq signal and MPRA activity (**Fig. S7C, Fig 4D**). ISM reveals the disruption of a p53 motif by the alternative allele (**Fig. S7C**). This variant is also an eQTL for *IFNAR2* in the lung (**Fig. S7D, Table 1**), supporting evidence that the variant perturbs a regulatory element important for *IFNAR2* expression in the lung. Finally, we highlight rs2297480 which causes gain of STARR-seq activity (**Fig. S7B**) and resides in the first intron of the main *Farnesyl Diphosphate Synthase* (*FDPS*) transcript where it is predicted to increase ATAC-and CAGE-seq signal, and *FDPS* expression (**Fig. S7E**). The alternative allele creates a G-rich motif positively contributing to the predictions (**Fig. S7E**). The variant is not an eQTL for *FDPS* in the lung but is an sQTL (**Fig. S7F, Table 1**). Though annotated as within the first intron of *FDPS*, in A549 cells rs2297480 overlaps a peak of H3K4me3 (**Fig. 2C**), indicative of promoter activity (**Fig. S7G, 2C**). Indeed, it is located in the promoter-proximal region of two *FDPS* transcript isoforms that exclude the penultimate exon of *FDPS,* and in the 5’ UTR of a third isoform (**Fig. S7H**). A CAGE-seq peak indicates the isoform(s) are being transcribed in lung (**Fig. S7H**). This suggests that the alt allele of rs2297480 may promote the transcription of an *FDPS* isoform missing the penultimate exon, that encodes part of the FDPS catalytic domain.

In summary, we demonstrated that for our data, both genomic and task-specific deep learning models, while having limited ability to identify amVars, can be used for hypothesis generation of observed and concordantly predicted variant effects.

## Discussion

### Using STARR-seq to identify functional single and combinatorial variants

In this study, we identify a set of severe COVID-19 associated risk variants which individually, and in some cases in combination, modulate regulatory activity in lung epithelial cells using a massively parallel episomal reporter assay – STARR-seq. Of 4,894 variants tested, only 29 modulated enhancer activity. This in line with a large-scale screen identifying only a small subset of variants as affecting regulatory elements, of which 90% were identified in only one out of two tested cell lines^37^.

For 8 of 49 risk loci, we identified multiple amVars, indicating GWAS associations may be driven by multiple regulatory modulating variants at a single locus, as previously proposed for melanoma and non-small cell lung cancer risk loci^16,38^. Consistent with this, 17.7% of eQTLs encompass multiple expression-modulating variants in strong LD^33^. This further emphasises the need to experimentally screen for functional risk variants, as statistical and predictive approaches cannot resolve clusters of variants in strong LD^32^. Multiple variants in LD across different enhancers at a locus may have regulatory function, in line with observations made in the context of obesity-associated variants at the *FTO* locus^39,40^. For Hirschsprung disease, three variants across different CREs were shown to synergistically reduce *RET* expression, amplifying the effect of individual variants^41^. This supports a model in which, rather than a single dominant causal variant, disease-associated loci can harbour multiple regulatory variants in LD, whose individually small effects compound to significantly influence gene expression. This could reflect cooperative interactions between multiple, individually weak, enhancers scattered across a locus^42^.

By testing variants in close proximity in combination we identified mainly additive, but also a few cases with apparent interdependent, effects on enhancer activity. This is consistent with a study finding non-additive effects of variant pairs residing within 150-bp using MPRA^43^. While one variant may be tolerated, multiple proximal variants may impair (or enhance) TF binding at a single or two adjacent TF motifs. We highlighted rs77819001 and rs76667867 as an example of apparent non-additivity but did not identify a common TF motif. This could be explained by the relevant motif not yet being known, which may be either a single or a composite motif comprising multiple TF binding sites that differ from the individual motifs^44^. In this scenario, only the combination of two variants would alter the DNA binding affinity sufficiently to cause loss of binding. Similarly, the flanking sequence is known to contribute to enhancer activity^45^, whereby alteration of a TF motif in combination with altered flanking sequence may exacerbate the effect of either case in isolation. Alternatively, these variants may reside in two independent TF binding sites for redundant activators or synergistic repressors.

### Deep learning models can complement STARR-seq data and aid hypothesis generation

We found that two recently developed deep learning models trained on large datasets had limited ability to accurately predict experimentally identified amVars. This is not surprising, as both the baseline activity in the STARR-seq assay and effect size of the amVars was relatively low, in line with observations from others^46^, leading to predictions where noise will have a large impact. Consistent with this, several other deep learning models showed limited accuracy in predicting allele-specific variants^33,47,48^. Instead of eliminating the need for experimental screens, particularly for variants with small effect sizes, deep learning models can complement high-throughput assays by contribution score attributions of experimentally determined variants and motif matching to generate hypotheses. This has been exemplified here, and by other recent application of deep learning models to explain observed regulatory _effects_34,48–51.

### Candidate loci and genes affecting severe COVID-19 outcomes

Several of the candidate target genes for variants identified here through STARR-seq in lung epithelial cells are in pathways with well-established links to COVID-19, lung inflammation, fibrosis and lung damage, and where known small molecule modulators could have therapeutic impact. STARR-seq can therefore be a valuable assay to investigate the functional effects, and potential direction of effect, of variants on target genes.

#### Interferon signalling

The largest allele-specific loss of STARR-seq activity we detected in A549 cells was at rs6517156 located in the last intron of *IFNAR2*. IFNAR2 is a type I interferon receptor that is key for immune responses to respiratory viruses, and the control of proinflammatory cytokines in the lung to protect against tissue damage post infection^52^. In humans, rare recessive *IFNAR2* loss-of-function variants result in increased risk of life-threatening respiratory infection^53^ and GWAS has shown that reduced *IFNAR2* expression is a risk factor for severe COVID-19 disease^2,3^. rs6517156 is also an eQTL for reduced *IFNAR2* expression in the lung, and is within an ENCODE CRE (EH38E3455612) that is a strong DNAse hypersensitive site in IMR-90 human fetal lung cells ^54^. Deep learning predicts the minor allele to decrease chromatin accessibility, and enhancer-associated histone modifications H3K27ac and H3K4me1, consistent with the rs6517156 alt allele disrupting an enhancer regulating *IFNAR2* expression in the lung. It also predicts that the alternative allele disrupts a binding motif for the transcription factor p53. In addition to genome stability and programmed cell death roles, p53 drives antiviral responses by regulating the expression of interferon response genes, while human coronavirus proteins promote proteasomal degradation of p53^55,56^.

STARR-seq also identified three rare variants (MAF<4%) at the *IFNA* locus on chromosome 9, that encodes a family of type I interferons. This locus evolved rapidly in mammals, presumably in response to pathogens^57^. All three rare variants result in reduced STARR-seq activity. Two of these rare variants are adjacent in the genome, lying in the intergenic region between *IFN10* and *IFN16* that appears to have derived from LINE-1 insertion during human/ape evolution. LINE-1 retrotransposons have been shown to contribute to the evolution of regulatory elements^58^.

#### Viral entry

On chromosome 1, rs2297480 at the *FDPS* locus is associated with increased STARR-seq activity. Though not an eQTL for *FDPS* in lung, the variant is an sQTL and marked by H3K4me3 in A549 cells. We propose the variant lies in a promoter-proximal enhancer or alternate promoter which produces an *FDPS* isoform that excludes the first and penultimate coding exon, missing part of the catalytic domain of farnesyl pyrophosphate synthase. FDPS is part of the mevalonate pathway and its reduced activity could impact protein prenylation, including that of Rab GTPases which control the endolysosomal pathways used by SARS-CoV-2 for cellular entry^59^. *RAB2A* is itself a risk locus for severe COVID-19^3^, and rs6471885, located 30kb upstream of the *RAB2A* promoter, is an eQTL for increased *RAB2A* expression in lung and showed increased STARR-seq activity in A549 cells.

#### Viral RNA degradation

The COVID-19-associated variant rs10774671, located at an intron/exon boundary, has been shown to affect *OAS1* splicing, causing isoform switching of *OAS1* to the enzymatically impaired p42 isoform^60–62^. *OAS1* is known to be important in sensing and degrading viral dsRNA, including SARS-CoV-2 RNA^63^. The rs4766668 variant, showing reduced STARR-seq activity, lies downstream of *OAS1* in the intergenic interval between *OAS1* and *OAS3*, and is an sQTL for *OAS1*.

#### Lung damage and repair

The alt allele at rs6133326, which reduced STARR-seq activity, is located 260kb upstream of *BMP2* in an ENCODE CRE showing strong DNAse accessibility and ATAC-seq signal in IMR90 lung cells. *BMP2* is up-regulated after epithelial injury and causes epithelial dysfunction and hyperpermeability^64^, and is down regulated in AT2 alveolar cells during early stages of lung regeneration^65^. Indeed, BMP2 signalling is thought to be important in pulmonary fibrosis^66^.

On chromosome 17, we prioritised five variant pairs at the *KANSL1* risk locus, encompassing the *CRHR1, MAPT* and *KANSL1* genes. Alt-alt alleles are associated with increased expression of a long non-coding RNA *LINC02210* and decreased *KANSL1* mRNA splicing in lung. Genetic variants at *CRHR1* have been associated with the response to corticosteroid treatment in asthma^67^, chronic obstructive pulmonary disease^68^, and in premature infants at risk for bronchopulmonary dysplasia^69^. Notably, dexamethasone, a corticosteroid, is the standard of care treatment for patients severely ill with COVID-19^70^. Similarly, variants at the *KANSL1* locus have previously been associated with lung fibrosis^71,72^, suggesting common underlying mechanisms. *MAPT* encodes the protein Tau, shown to aggregate in brain cells following cleavage by SARS-CoV-2 3CL proteases^73^.

By integrating STARR-seq and deep learning models, we identify functional variants affecting regulatory activity in isolation and in combination in the lung epithelium. We propose mechanisms by which these variants may influence COVID-19 pathogenesis, thereby laying the groundwork for follow-up investigations. We propose that the variants we identified in our STARR-seq screen are highly suitable for endogenous validation and follow-up studies, for example using prime editing to generate homozygous risk alleles to identify the target gene(s) and effect on response to viral infection.

Our data has several limitations. Firstly, our screen was limited to one lung epithelial cell line, therefore, variants exerting their effect in a different cell type, primarily immune but also other lung cell types, are not captured. A substantial proportion of our identified allele-specific variants resides within inaccessible chromatin regions in A549 cells, but may be endogenously functional in other cell types. A subset of amVars that showed gain-of-activity, however, may drive *de novo* chromatin accessibility by encoding favourable motifs for TF binding. Secondly, we assessed variants under homeostasis. IL-1ß treatment and SARS-CoV-2 infection, for example, have previously revealed context-dependent variant effects^7,74^, suggesting additional amVars would be identified in the inflammatory state caused by viral infection. Regardless, assays in cell models are always likely to miss some variants that are functional *in vivo.* Thirdly, STARR-seq is an episomal assay, lacking chromatin context, and may be confounded by mRNA stability and splicing effects given the self-transcribing design^75^. Further, we assessed the ability of AlphaGenome and Malinois to predict amVars, however, we did not perform a systematic analysis of available deep learning models for variant prediction, and only had a limited set of amVars for comparison. Lastly, our study was limited to the prioritization of variants with regulatory effects, but endogenous validation of variant effects and target gene identification will be required.

## Methods

### Variant selection and library design

The 95% credible set from the second GenOMICC release^2^, containing causal variants to 95% statistical probability, and the GWAS summary statistics for the third GenOMICC release ^3^ were kindly shared by the authors. For the third release, variants adding up to a cumulative posterior inclusion probability of 99% for each risk locus were included in the STARR-seq library. Variants in LD with any of the 49 lead variants from the third GenOMICC release were extracted using the *LDproxy_batch* function from *LDlinkR* v.1.2.1 using the GRCh38 genome build^76^. The resulting 99% credible set from^3^, the 95% credible set from^2^, and variants in LD were merged and duplicates removed, giving 4,894 unique variants.

An oligonucleotide pool consisting of 170-bp genomic sequence centred on each of the 4,894 variants, as both reference and risk allele, was designed using *snp2fasta* (https://github.com/efriman/snp2fasta) with the parameters *--flank 85 --combinations 5 --maxdistance 100*. This generated additional oligonucleotides for up to 5 variants occurring within 100 bp genomic distance as all possible combinations of reference and alternate allele.

As positive controls, we included 61x 170-bp sequences centred on the highest STARR-seq peaks in A549 lung adenocarcinoma cells (untreated condition, 0h) by sorting the bigwig signal in peaks from^27^ (NCBI GEO accession: GSE114063). We included 119x putative active sequences centred on the highest H3K27ac peaks in the H358 bronchioalveolar carcinoma cell line (NCBI GEO_GSM1635574), whereby each sequence required at least 30% overlap with ENCODE cCREs downloaded from https://api.wenglab.org/screen_v13/fdownloads/cCREs/GRCh38-ELS.bed (note that these are not used in this study and were excluded from the results presented). As negative controls, we included 10 scrambled sequences of A549 positive controls, 30 scrambled sequences of H358 putative active sequences, and 10 x 170 bp genomic sequences devoid of A549 chromatin modifications (A549 ENCODE H3K4me1, HeK4me3, H3K27ac, H3K9me3, ATAC-seq), or A549 STARR-seq enrichment from^27^. Lastly, we included 8x primer-amplified A549 controls from a previous test run of only controls sequences (**Table S7**). This resulted in 9,788 single variant oligonucleotides, 3776 combinatorial oligonucleotides and 238 controls (**Table S1**). 15-bp flanking adapters with primer binding sites for PCR amplification were added (FW: ACGCTCTTCCGATCT, RV: GTGCTCTTCCGATCT) and the resulting STARR-seq library consisting of 13,802x 200-bp sequences synthesised as pooled oligonucleotides from Twist BioSciences (USA).

### STARR-seq plasmid library construction

The STARR-seq plasmid library was generated according to the UMI-STARR-seq protocol^28^ with minimal alterations. Briefly, the protocol involved candidate sequence PCR amplification, digestion of the hSTARR screening vector (Addgene #99296), cloning of library inserts into the digested hSTARR vector and plasmid library amplification and purification. Diverging from the UMI-STARR-seq protocol, the oligonucleotide pool was amplified using the following PCR programme: 95°C for 3min, followed by 14 cycles of 98°C for 20s, 65°C for 15s and 72°C for 15s, and a final step at 72°C for 1min. The optimal number of cycles was determined by a qPCR test reaction using the same conditions, except 0.25 μl Evergreen dye (Biotium #31000-T) was added to the reaction and 30 cycles performed followed by a plate read step. Amplified oligonucleotides were purified using the QIAquick PCR purification kit (Qiagen #28104) according to manufacturer’s instructions while omitting the AMPureXP bead size-selection of amplified libraries due to the fixed length of the oligonucleotides. Purified library inserts were cloned into the digested hSTARR vector using NEBuilder HiFi DNA Assemble In-Fusion HD (NEB #E2621L) in 2 reactions (each reaction: 100 ng digested hSTARR plasmid, 2x molar excess library insert, 5 μl NEBuilder reaction mix, to 10 μl with H_2_O) by incubation in a thermocycler for 15 min at 50°C. Following transformation of the resulting STARR-seq library according to the UMI-STARR-seq protocol^28^, the library was purified using four columns of the Qiagen Maxiprep Plus Kit (Qiagen #12963) according to manufacturer’s instructions.

### STARR-seq screen in A549 cells

A549 lung adenocarcinoma cells (ATCC #CLL-185) were maintained in DMEM (Life Technologies #41965039) supplemented with 10% foetal calf serum (FCS) and 1% penicillin/streptomycin at 37°C with 5% CO_2_ and passaged every 2-3 days. For each of five biological replicates, performed on different days, 4×10^7^ cells were resuspended in 375 μl electroporation buffer (MaxCyte #EPB1) and electroporated with 80 μg STARR-seq plasmid library (see above) making up 25 μl (400 μl total) in an OC-400 electroporation cuvette (MaxCyte #GOC4) using the manufacturer-pre-set ‘A549’ protocol on a MaxCyte GTx system. Following electroporation, cells were let to recover for 25 min at 37°C before gentle transfer to a culture flask with pre-warmed DMEM supplemented with 10% FCS but without antibiotics. Electroporated cells were lysed and total RNA harvested after 6h, and output libraries processed according to the published UMI-STARR-seq protocol ^28^ with minimal alterations. Briefly, this involved mRNA isolation, reverse transcription of reporter transcripts, purification, and PCR amplification. Diverging from the UMI-STARR-seq protocol, upon unique molecular identifier (UMI) introduction by PCR, a modified primer additionally introducing an i7 index (CAAGCAGAAGACGGCATACGAGATNNNNNNNNNN[i7]GTGACTGGAGTTCAGAC GTGT*G) was used instead of the P7-UMI primer, allowing for dual indexing of sequencing libraries in combination with the Illumina i5 indexing primer (NEB #7600). Output sequencing libraries were prepared according to the UMI-STARR-seq protocol except performing eight PCR reactions, using only 5 μl junction PCR product per reaction and amplifying for 25 PCR cycles followed by library purification using AMPure XP beads using 0.9 vol beads to 1 vol output sequencing library.

Input sequencing libraries were prepared directly from the STARR-seq plasmid library in duplicate according to the UMI-STARR-seq protocol except introducing both i5 and i7 dual indexes and 10-bp UMIs as for the output sequencing libraries (see above). Input and output libraries were sequenced (2×150 paired-end) on a NextSeq2000 (Illumina #SY-415-1002) using the NextSeq 1000/2000 P1 Reagents Kit (300 cycles) (Illumina #20050264). The index 1 read was extended to 18 cycles to sequence both the index and the UMI. Per replicate, between 24-46 million sequencing reads were obtained.

### STARR-seq data processing and analysis

Sequencing reads were aligned to a custom reference genome containing the STARR-seq library oligonucleotide sequences using bowtie1 v.1.3.0^77^ with the parameters *-X 300-p 4-v 3 --norc-m 1 --best --strata* to allow for 3 mismatches and unique alignment only. Aligned sam files were converted to bam file format using samtools v.1.6^78^ and paired end reads aligning to the same sequence featuring identical UMIs collapsed using the umi_tools dedup function from UMI-tools v.1.1.4^79^ with the *--paired* option, thereby removing PCR duplicates. Counts for each library sequence were generated from aligned, UMI-collapsed reads using the htseq-count function with the parameters*-s no-m intersection-strict-a 30-f bam* from the HTSeq package v.2.0.2^80^. We required sequences to have at least 10 UMI-collapsed read counts for each of the two input replicates and at least 5 UMI-collapsed read counts for each of the five output replicates. The STARR-seq activity for each remaining library sequence (13,461/13,802 oligonucleotide sequences) as log2FC of output/input libraries as well as normalised read counts were computed using DESeq2 v.1.44.0^81^ in R v.4.4.0 using default parameters with the design formular *∼type* and *fitType=”local”* (**Table S2**). Library sequences with a log2FC greater than 1 at an adjusted *p*-value of <0.01 were considered active.

Allele-specific activity was assessed using the mpralm R package^82^, a linear model developed for MPRA data based on the voom framework^83^, with the parameters *normalize=TRUE*, *aggregate=“none”, block=block_vector, model_type=“corr_groups”, plot=TRUE* to estimate log2FC between alleles and test for differential activity using moderated t-statistics (**Table S3**). Variants were considered to be amVars if the mpralm adjusted p-value was < 0.01 and at least one of reference and risk allele were determined to be active by DESeq2 analysis (see above). For combinatorial variant pair oligos we performed pairwise contrast testing between the four alleles and computed significance using the *mpralm “eBayes”* function (**Table S4**).

Variants were annotated by their genomic location using *annotatePeaks.pI* program from HOMER tools v4.1 with default parameters ^84^, providing the hg38 human gene transcript reference file as *-gtf*, downloaded from UCSC (hgdownload.soe.ucsc.edu/goldenPath/hg38/bigZips/genes/hg38.refGene.gtf.gz).

### Intersection with A549 chromatin datasets ENCODE cCREs

A549 datasets indicative of endogenous enhancer function were downloaded as narrow_peak files from ENCODE, including for ATAC-seq (ENCFF648AEN), DNase-seq (ENCFF128ZVL), H3K4me1 ChIP-seq (ENCFF594YDK), H3K4me3 ChIP-seq (ENCFF404REU) and H3K27ac ChIP-seq (ENCFF747IZX). Variant coordinates were intersected with peak coordinates using the *intersect* function from BEDTools v2.27.1, providing variants as-a and A549 datasets as-b. ENCODE A549 candidate CREs were downloaded from ENCFF767VHY and intersected with STARR-seq library oligonucleotide coordinates (170bp) using BEDTools intersect.

### Computing the interaction of variant combinations

To calculate the expected STARR-seq log2FC in presence of both variants (alt_alt) based on the observed STARR-seq activity in presence of either variant alone (ref_alt and alt_ref), we computed the expected log2FC for the alt_alt oligonucleotide if the effect of both variants combined in an additive or in a multiplicative manner (**Table S5**) as follows:

Expected _additive_ = log_2_(2^A^ + 2^B^ - 2^C^)

Expected _multiplicative_ = A + B - C

A, B and C are the observed log2FC for ref_alt allele (A), alt_ref allele (B) and ref_ref allele (C), respectively. For one variant combination, the expected additive log2FC was set to 0 as the theoretically expected fold-change of RNA/DNA was negative and negative log2FC corresponds to no STARR-seq activity, hence a negative log2FC and a log2FC of 0 represent the same outcome.

### Predicting variant effects with GTEx, AlphaGenome and Malinois

The effect of variants on gene expression (eQTLs) and mRNA splicing (sQTLs) in lung tissues was obtained from querying the GTEx Portal (https://gtexportal.org/home/).

We predicted the impact of amVars and non-amVars within STARR-seq active sequences on ATAC-seq, DNase-seq, CAGE-seq, RNA-seq and ChIP-seq signals (TF binding and histone modifications, including H3K27ac, H3K4me1, H3K4me3) in A549 cells using AlphaGenome v.0.1.0^34^. We used the predict_variant dna model with default parameters, considering a sequence interval of 1 Mb centred on the variant. Given AlphaGenome returns low-confidence predictions even if no true change is predicted, we filtered for high-confidence predictions using the quantile score (absolute value>0.99) (**Table S6**), which represents the predictions rank within a background distribution of GnomAD common variants for the predicted feature^34^. A prediction with a quantile score of 0.99 is within the 99^th^ percentile of common variant predictions for the features assessed (e.g. ATAC-seq). Genome tracks show the difference in prediction between alternative and reference alleles (alt – ref).

To compute the contribution of each nucleotide to a predicted feature within a 170-bp window centred on the variant, reflecting the STARR-seq oligonucleotide length, we performed *in-silico* mutagenesis (ISM) using the *score_ism_variants* model and visualised scores using the *plot_components* function.

For predicting A459 MPRA activity using Malinois, the STARR-seq libray was filtered for entries of 170 bp (i.e. excluding insertions/deletions). Sequence padding, activity predictions, and contribution score calculations were done using procedures described in https://github.com/sjgosai/boda2 using the available Malinois model trained on A549 cells ^35^.

### Motif analysis

Contribution scores from Malinois were used to find sequence patterns using the modisco-lite 2.3.2 *motifs* command with *-n 50000*^85^. Fi-NeMO (https://github.com/kundajelab/Fi-NeMo) *call-hits* was used to determine the genomic coordinates of pattern matches.

## Data availability

Raw STARR-seq data and processed files generated from this study have been deposited in the Genome expression Omnibus (GEO) repository under the accession number GSE320469. Publicly available A549 datasets were downloaded from ENCODE (https://www.encodeproject.org), including ATAC-seq (ENCFF648AEN), DNase-seq (ENCFF128ZVL), H3K4me1 (ENCFF594YDK), H3K4me3 (ENCFF404REU) and H3K27ac (ENCFF747IZX). The Malinois A549 model was downloaded from https://zenodo.org/records/10698014.

## Code availability

The SNP2fasta package generated to obtain fasta files for single and combinatorial variant oligonucleotide libraries is available on GitHub: https://github.com/efriman/snp2fasta

## Author contributions

G.W., E.T.F, W.A.B. and S.C.B. designed this study. G.W. performed STARR-seq experiments and data analyses, the latter together with E.T.F. G.W. generated visualizations and wrote the manuscript which was edited by E.T.F, W.A.B. and S.C.B.

## Acknowledgements

We thank Erola Pairo-Castineira, Konrad Rawlik, J. Kenneth Baillie for sharing GWAS fine-mapping results and discussion on variant selection for the purpose of library design. We also thank Veronique Vitart for discussion on variant selection and Luciana Gómez-Acuña for advice on experimental techniques. Sequencing was performed at the Genetics Core of the Edinburgh Clinical Research Facility. We thank the Institute of Genetics and Cancer core FACs and technical services facilities for their support. This work has made use of the resources provided by the Edinburgh Compute and Data Facility (ECDF) (http://www.ecdf.ed.ac.uk/).

---

**Fig S1.**
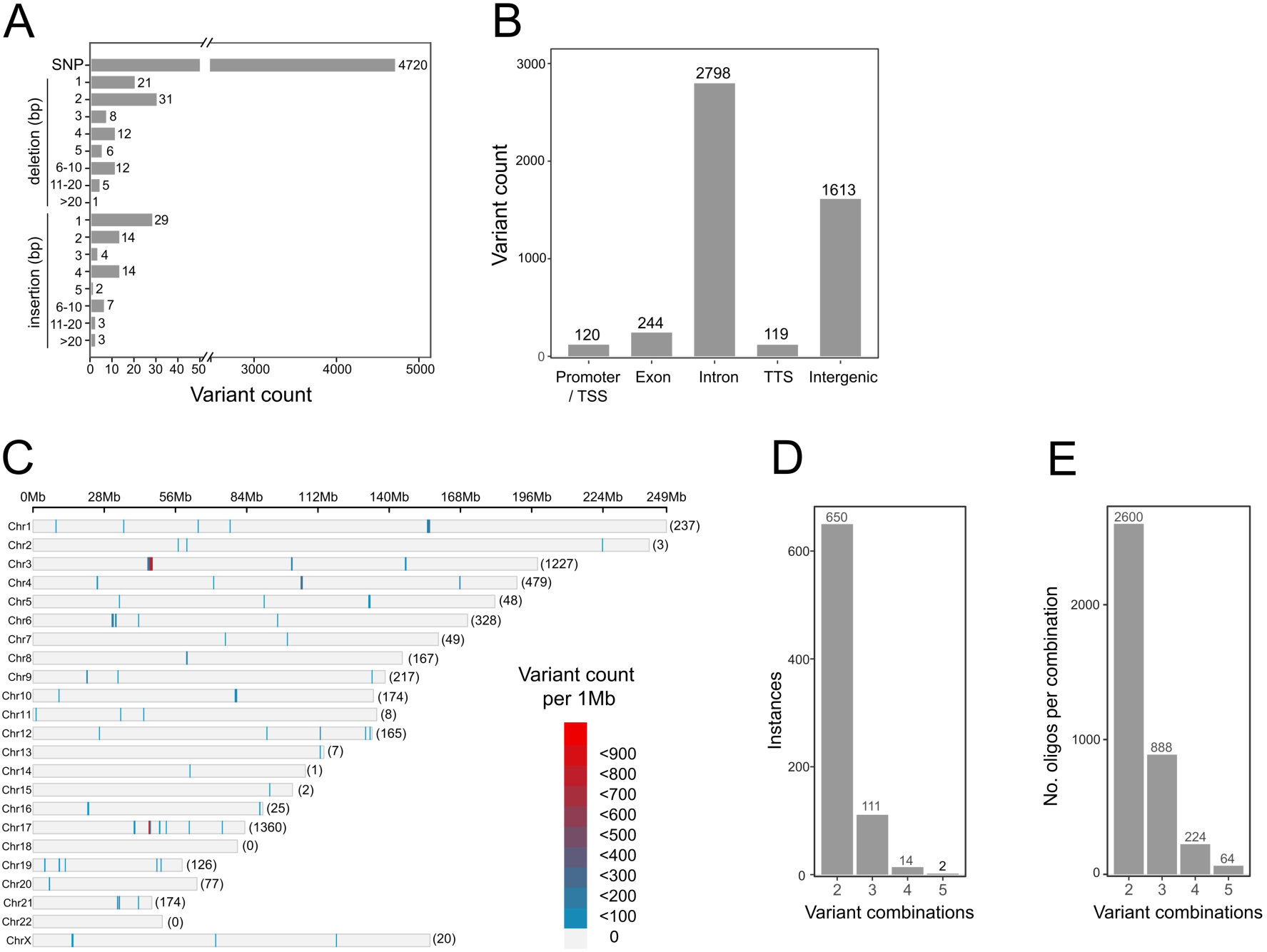
STARR-seq library variants are primarily non-coding SNPs concentrated at a few risk loci. (A) Number of STARR-seq library variants by mutation type; single nucleotide polymorphisms (SNPs), and small insertions and deletions. (B) Classifications of STARR-seq library variants by genomic location; promoter/transcription start sites (TSS), exons, introns, transcription termination sites (TTS) and intergenic regions. (C) Density of STARR-seq library variants per megabase (Mb) for each chromosome. Number of STARR-seq library variants per chromosome in parentheses. (D) Instances of 2, 3, 4, and 5 variants occurring within 100-bp which were included as combinations in the STARR-seq library. (E) Number of combinatorial oligonucleotides included in the STARR-seq library for combinations of two to five variants as shown in (D).

**Fig S2.**
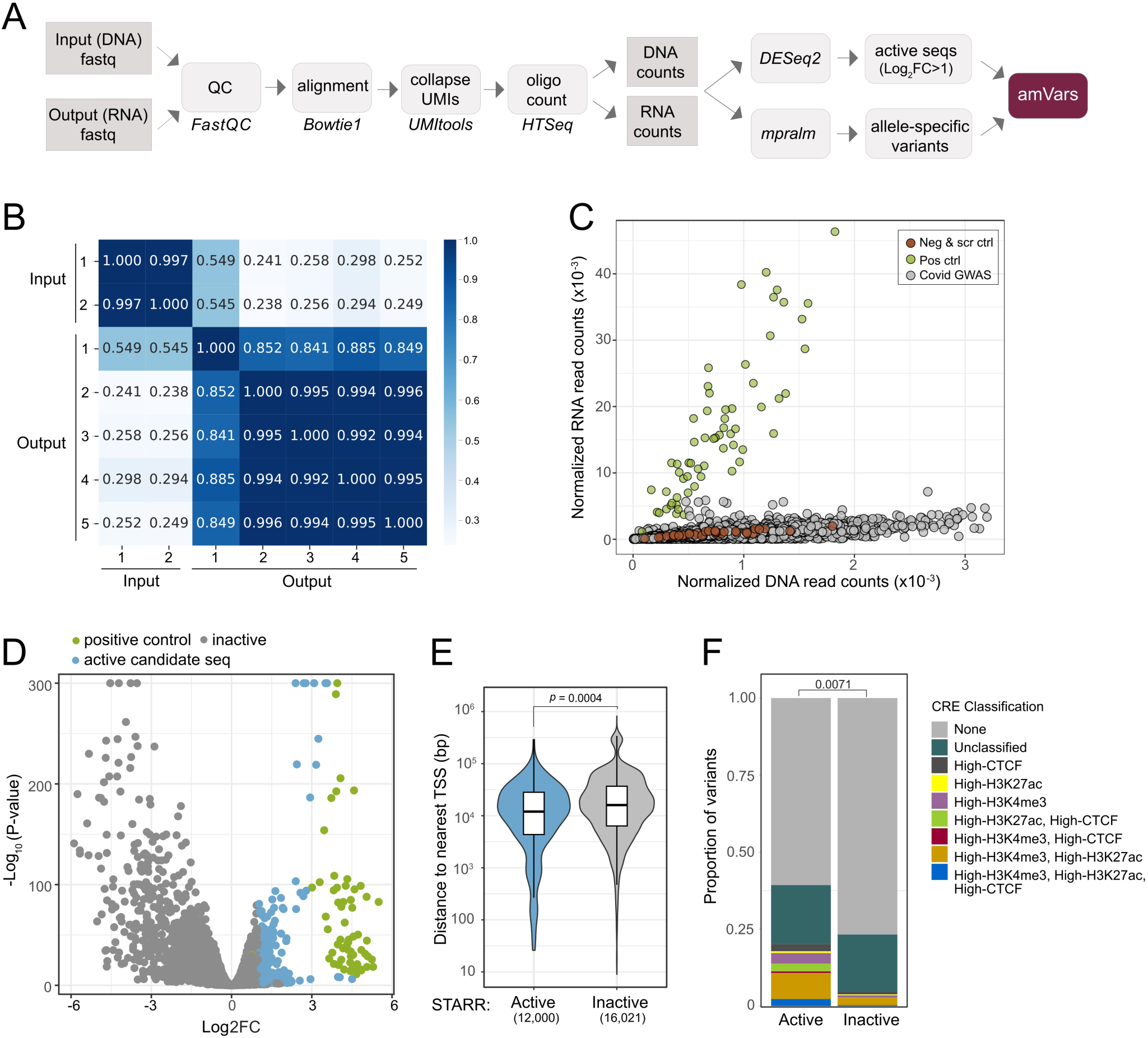
Properties of STARR-seq active sequences. Related to Fig. 2 (A) Schematic of STARR-seq analysis workflow. Input (DNA) and output (RNA) sequencing fastq files for each replicate were assessed for sequencing quality using *FastQC* and aligned to a custom reference genome containing STARR-seq library sequences using *Bowtie1.* Reads aligning to the same sequence with identical unique molecular identifiers (UMIs) were collapsed using *UMItools* and read counts for each library sequence generated using *HTSeq.* STARR-seq activity was computed using *DESeq2*, considering sequences as active at a log2FC > 1 (FDR < 0.01), and allelic differences between reference and risk alleles assessed using *mpralm* (FDR < 0.01). (B) Pearson correlation of input and output reads for all replicates. (C) *DESeq2* normalised read counts in RNA (output) against DNA (input) libraries, averaged across biological replicates, coloured by category. (D) Volcano plot showing *DESeq2*-computed significance as-log_10_(*P-*value) adjusted for multiple testing against log2FC (STARR-seq activity) for all sequences tested. (E) Distance to the nearest TSS (in bp) for STARR-seq active and inactive sequences. Boxplot showing the second quartile (upper half), median (middle line) and second quartile (lower half). (F) Proportion of STARR-seq active and inactive sequences overlapping A549 ENCODE candidate *cis-*regulatory elements (cCREs). As not all datasets are for A549 were used by ENCODE for classification, lacking DNase-seq data, cCREs with low H3K27ac, H3K4me3 or CTCF were designated unclassified by ENCODE. Sequences classed as inactive by ENCODE are displayed as None. Differences in CRE distribution were assessed using Fisher’s exact test.

**Fig S3.**
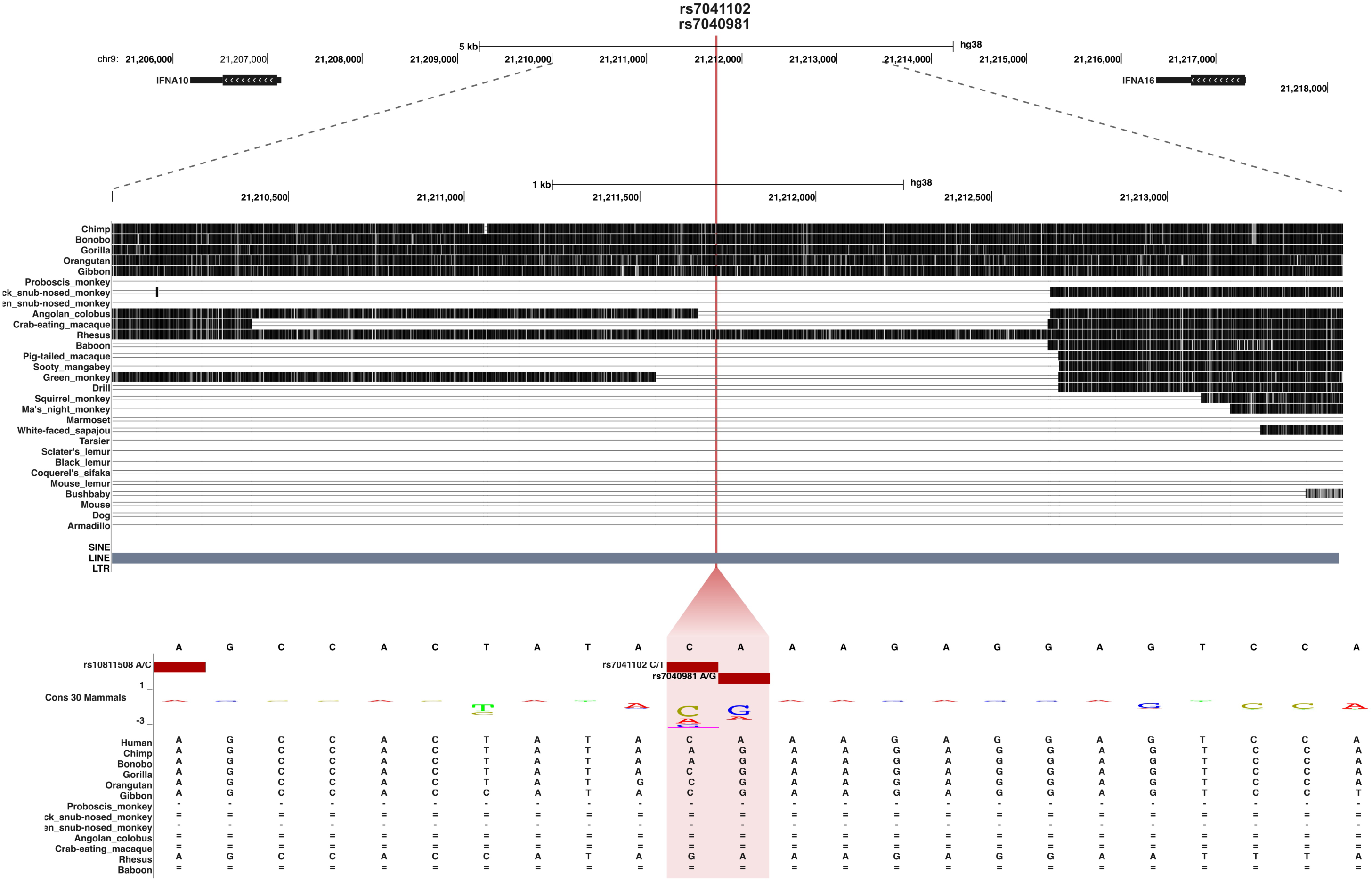
Adjacent rare variants with differential STARR-seq activity are located in a recent LINE insertion. UCSC genome browser screen shot (GRCh38) showing SNPs rs7041102 and rs7040981 located upstream of *IFNA10* and downstream of *IFNA16*, two genes encoding type I interferon alphas. Sequence conservation to primates and other mammals, and the position of a LINE-1 insertion specific to humans and apes is shown below.

**Fig S4.**
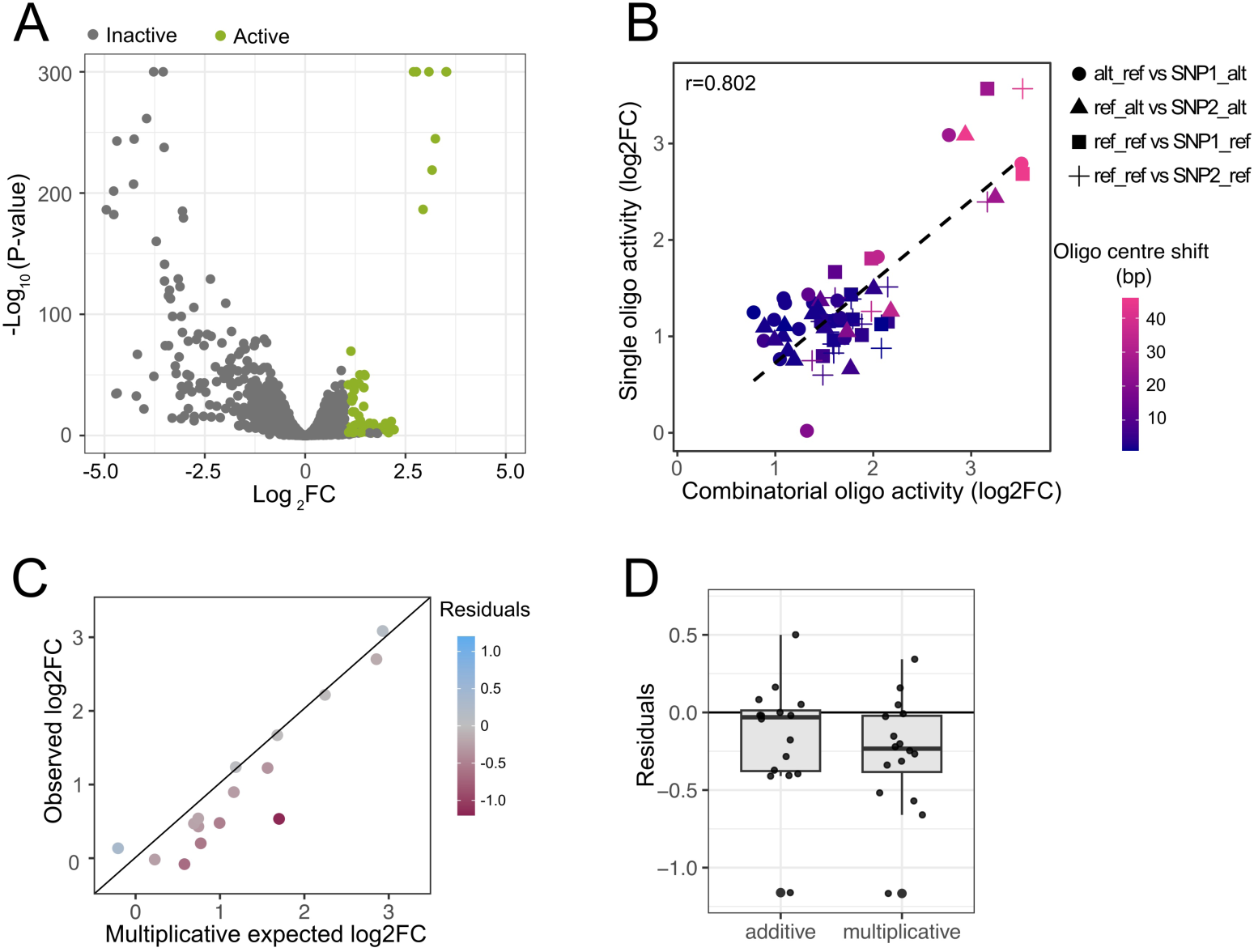
Characterization of combinatorial variants. (A) Volcano plot showing STARR-seq-log10(*p*-value) adjusted for multiple testing against activity as log2FC computed using *DESeq2* for 3658 combinatorial variant oligonucleotide candidate sequences that passed filtering. Active sequences (log2FC >1, FDR < 0.01) highlighted in green. (B) Correlation between STARR-seq activity of variants tested as single variant-and combinatorial-oligos matched in genotype. Reference (ref_ref) combinatorial allele compared to single variant reference alleles for variant one and variant two alone, ref_alt combinatorial allele compared to variant 2 alternative allele alone, and alt_ref combinatorial allele compared to variant 1 alternative allele alone. As combinatorial oligonucleotides are centred on the middle between variants while single variant oligos are centred on the variant, the colour indicates the shift in oligonucleotide centre. (C) STARR-seq observed Δlog2FC (log2FC alt - log2FC ref) against the theoretically expected Δlog2FC for alt_alt alleles if both variant effects interacted multiplicatively. Colour indicates the difference between observed and expected effects (Residuals). (D) Difference between observed and expected log2FC (residuals) for alt_alt alleles for additive and multiplicative interaction terms. Negative values indicated greater-than-expected loss of activity based on either variant alone. Each point indicates a variant pair. Lower and upper hinges of the boxplot correspond to the first and third quartile, respectively, the middle representing the median.

**Fig S5.**
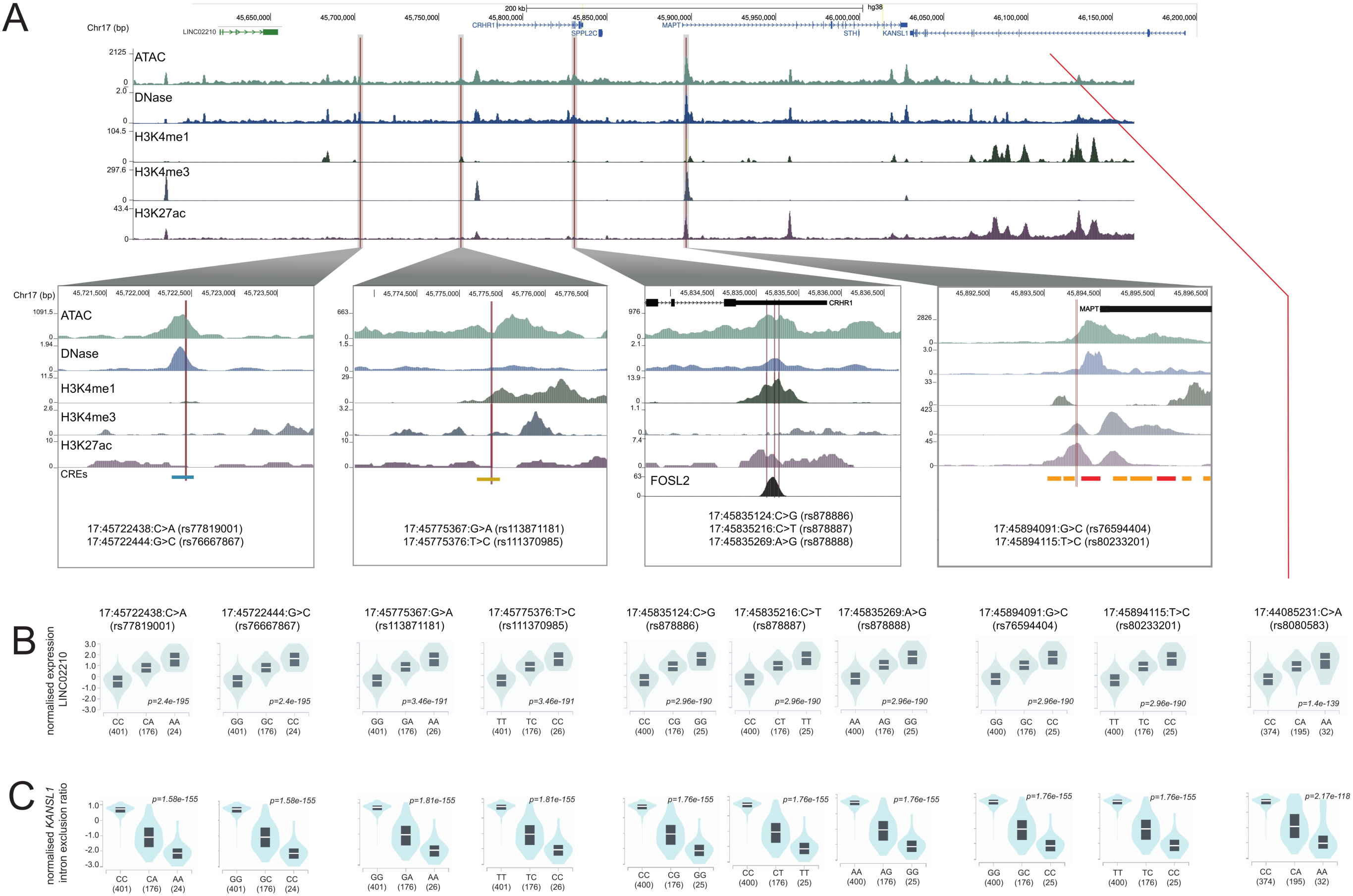
Chromatin features of prioritised variant pairs at the *CRHR1*-*KANSL* locus. (A) UCSC Genome Browser view (hg38) of the *CRHR1-KANSL* GWAS risk locus featuring five prioritised variant combinations, showing overlap with ENCODE ATAC-seq, DNase-seq, and H3K4me1, H3K4me3, H3K27ac ChIP-seq data from A549 cells and ENCODE cCREs. Below; Zoom-ins of 5-kb centred on variant pair, additionally showing overlap with ENCODE FOSL2 ChIP-seq density. (B) GTex eQTL violin plots showing the normalized expression of long non-coding RNA *LINC02210* in lung tissue for each genotype of variants displayed in (A). (C) GTex sQTL violin plots showing the normalized KANSL1 chr17:46094701:46170855 intron exclusion ratio in lung tissue for each genotype of variants displayed in (A).

**Fig S6.**
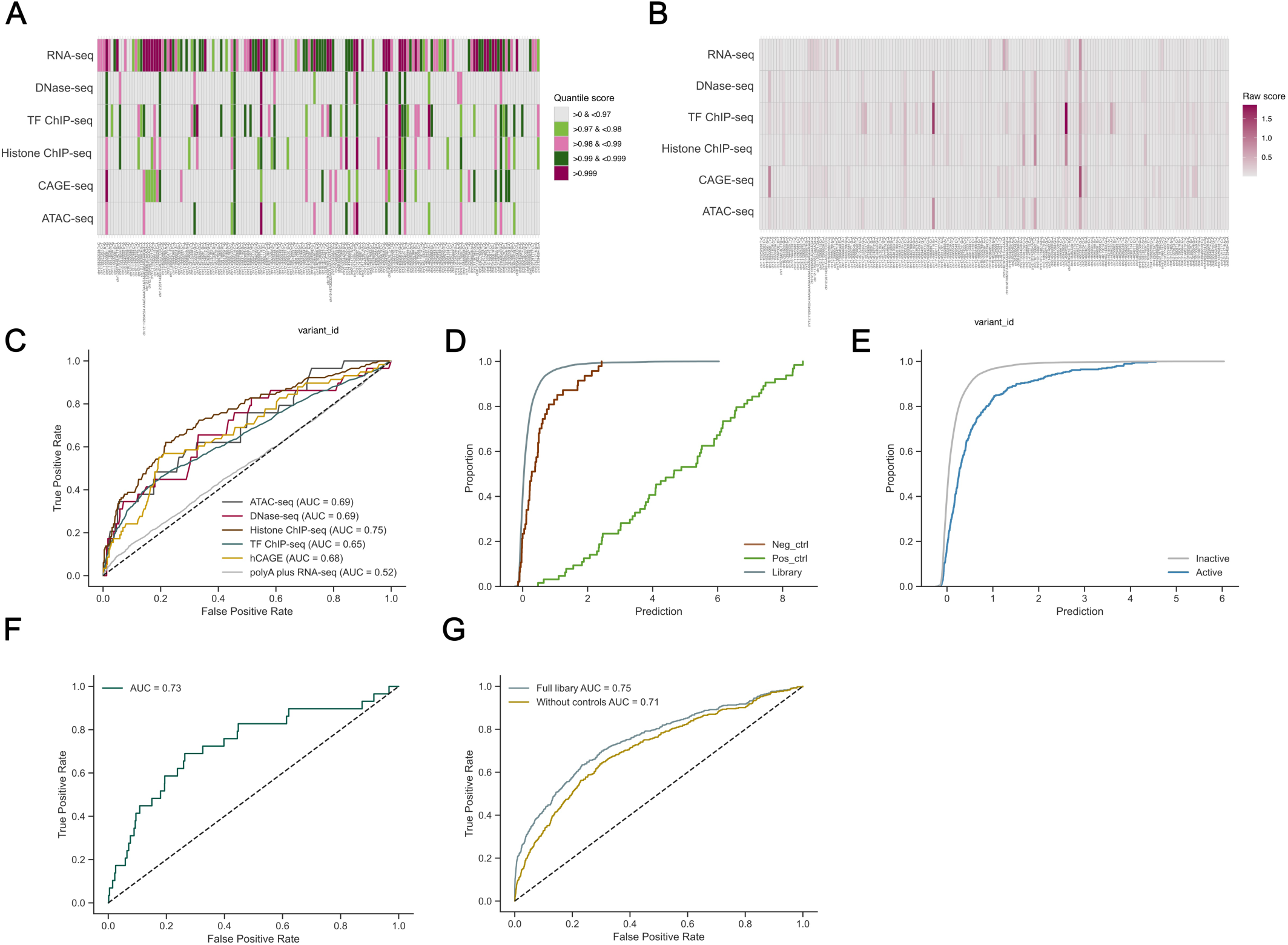
Comparison of AlphaGenome and Malinois predictions for genomic features and MPRA activity and comparison to STARR-seq (A) Heatmap showing AlphaGenome absolute quantile scores for predictions for 166 STARR-seq active variants across six features, including RNA-seq, DNase-seq, ChIP-seq for transcription factors, ChIP-seq for histone modifications (includes H3K27ac, H3K4me1, H3K4me2, H3K4me3), CAGE-seq and ATAC-seq. Colour indicates the quantile score, grouped into bins. Where multiple predictions of the same feature for one variant were generated (e.g. for TF and histone ChIP-seq and RNA-seq), the highest absolute quantile score was considered. (B) Heatmap of AlphaGenome absolute raw scores (effect sizes) for 166 STARR-seq active variants predicted across six features as in (A). (C) Receiver operating characteristic (ROC) curve for AlphaGenome predictions of variants effects (i.e. loss, no effect, or gain), separated by feature. (D) Cumulative proportion of Malinois prediction scores for STARR-seq categories. (E) Cumulative proportion of Malinois prediction scores for active and inactive STARR-seq sequences from the COVID-19 variant library. (F) ROC curve for Malinois A549 MPRA activity predictions of variant effects (i.e. loss, no effect, or gain). (G) ROC curve for Malinois A549 MPRA activity predictions of STARR-seq activity (i.e. active/inactive) for the whole library or excluding positive and negative controls.

**Fig S7.**
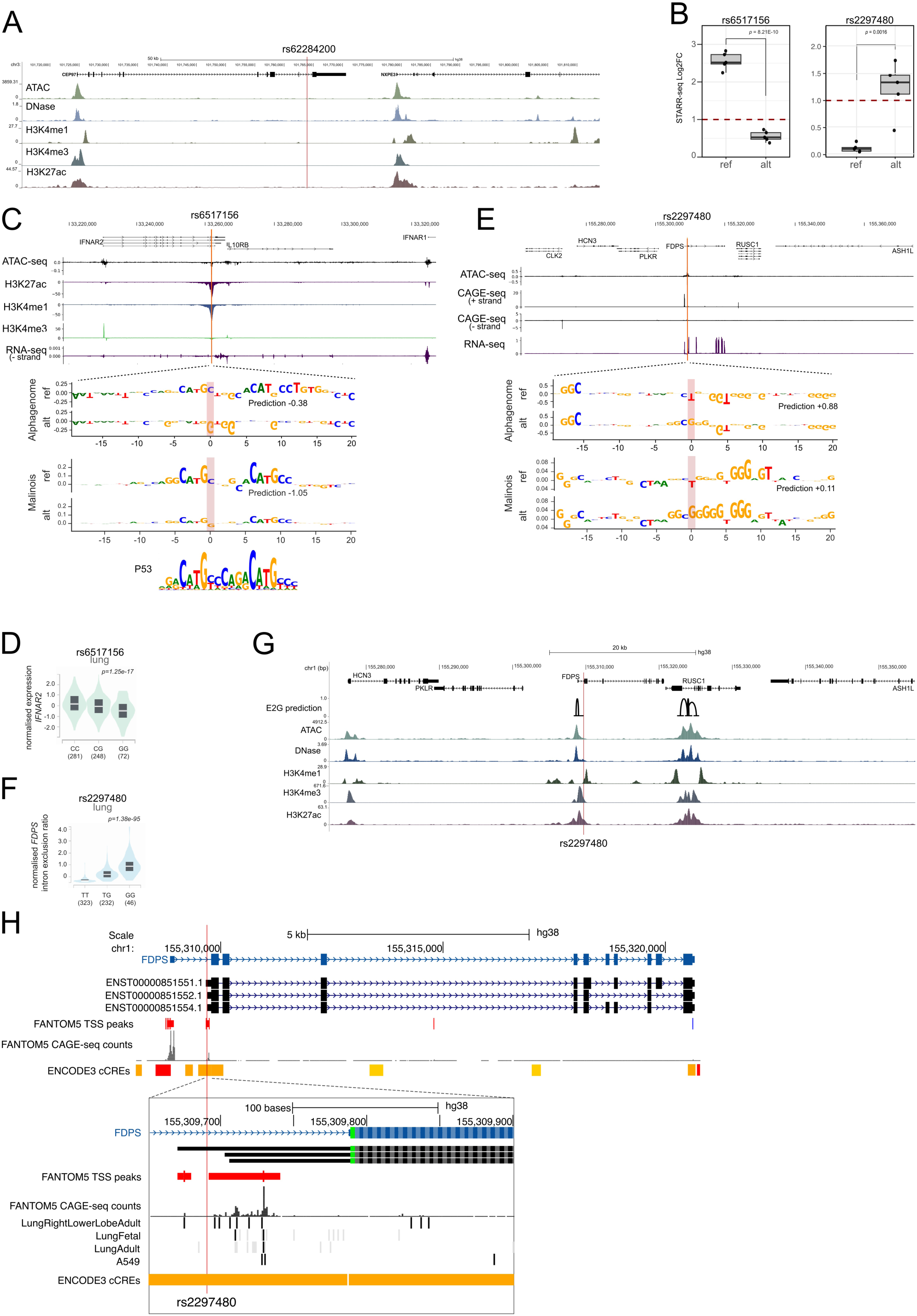
*In-silico* mutagenesis of candidate enhancers. (A) UCSC genome browser tracks (hg38) centred on rs62284200, showing A549 ATAC-seq, DNase-seq and ChIP-seq for H3K4me1, H3K4me2, H3K4me3 and H3K27ac datasets from ENCODE. (B) STARR-seq activity of (left) rs6517156 and (right) rs2297480 for reference and alternative alleles. Boxplot showing the second quartile (upper half), median (middle line) and second quartile (lower half), and five biological replicates displayed as individual points. Dotted red line indicates the STARR-seq activity threshold (log2FC > 1). (C) Top; AlphaGenome alternative-reference predicted genome tracks for rs6517156 centred on the variant in A549 cell, showing the difference in predictions between alternative and reference allele for features with an absolute quantile score > 0.99. Below; AlphaGenome (top panel) and Malinois (bottom panel) contribution scores for reference and alternative alleles of rs6517156 for ATAC-seq and MPRA activity predictions, respectively. The matched HOCOMOCO motif for p53 (P53_HUMAN.H11MO.0.A) is at the bottom. (D) GTex eQTL violin plots showing effect of ref and alt alleles at rs6517156 on *IFNAR2* expression in lung tissue. (E) As in (C) but for rs2297480 and showing AlphaGenome and Malinois comparative *in-silico* mutagenesis for reference and alternative alleles showing the contribution scores to CAGE-seq and MPRA activity predictions, respectively. (F) GTex sQTL violin plots showing the normalized *FDPS* chr1:155319688-155320409 intron exclusion in lung tissue. (G) UCSC genome browser tracks centred on rs2297480, showing A549 ATAC-seq, DNase-seq and ChIP-seq for H3K4me1, H3K4me3, H3K27ac datasets from ENCODE. ENCODE Enhancer2Gene (E2G) predictions for A549 cells showing rs2297480 is predicted to reside in an enhancer regulating *FDPS*. (H) UCSC genome browser tracks showing the reference (blue) and three alternative transcript isoforms excluding the first non-coding exon (black) of *FDPS*, FANTOM5 TSS peaks and total CAGE-seq counts as well as ENCODE cCREs. Zoom-in: Additional FANTOM5 CAGE-seq reads in selected lung tissue samples and A549, showing that rs2297480 resides near or at an alternative promoter active in lung.

**Table S1. COVID-19 STARR-seq library sequences.** Library sequences in fasta format, including single, combinatorial and control sequences. 15bp adapters on either end in upper case.

**Table S2. STARR-seq results for all library sequences.** Showing candidate sequence activity as log2FC with padj as computed using *DESeq2*, including normalized input and output read counts averaged across biological replicates.

**Table S3. STARR-seq results for single variant candidate sequences.** Including information on STARR-seq activity as determined using *DESeq2* and allele-specific effects determined using *mpralm*, read counts, variant coordinates, GC content, result and fasta sequence.

**Table S4. Allele-specific activity for 16 active variant pairs.** *mpralm* computed pairwise comparisons for all alleles of 16 variant combinations where at least one allele showed log2FC>1.5.

**Table S5. Observed and expected combinatorial variant effects under an additive and multiplicative model.** (Columns B-E) Observed STARR-seq activity as log2FC for all allelic combinations 16 variant pairs where at least one allele showed STARR-seq activity (log2FC>1.5, FDR<0.01). Expected log2FC for the alt_alt allele under an additive (column F) and multiplicative (column G) model, and discrepancy between predicted and observed STARR-seq log2FC (columns H-I).

**Table S6. AlphaGenome predictions of variant effects on A549 genomic features for amVars and non-amVars within STARR-seq active sequences.** AlphaGenome raw score indicates the magnitude of predicted change, the quantile score represents the predictions’ rank within a background distribution of GnomAD common variants for the predicted feature (here filtered for predictions >0.99), and assay title identifies the feature predicted. The predicted target gene (for RNA-seq), TF (for TF ChIP-seq) or histone modification (for histone ChIP-seq) is indicated. Δlog2FC_mpralm represents the change in STARR-seq activity caused by the alternative allele (alt-ref).

**Table S7. Primer sequences used to amplify eight sequences during a previous test run.**

